# Data based slurry treatment decision tree to minimise antibiotic resistance and pathogen transfer while maximising nutrient recycling

**DOI:** 10.1101/2022.02.25.481976

**Authors:** Thi Thuy Do, Stephen Nolan, Nicky Hayes, Vincent O’Flaherty, Catherine Burgess, Fiona Brennan, Fiona Walsh

**Author notes:** Corresponding author: Fiona Walsh, Thi Thuy Do **Email:**.

## Abstract

Direct application of pig slurry to agricultural land, as a means of nutrient recycling, introduces pathogens, antibiotic resistant bacteria, or genes, to the environment. With global environmental sustainability policies mandating a reduction in synthetic fertilisation and a commitment to a circular economy it is imperative to find effective on-farm treatments of slurry that maximises its fertilisation value and minimises risk to health and the environment. We assessed and compared the effect of storage, composting, and anaerobic digestion on pig slurry microbiome, resistome and nutrient content. Shotgun metagenomic sequencing and HT-qPCR arrays were implemented to understand the dynamics across the treatments. Our results identified that each of the treatment methods had advantages and disadvantages, depending on the parameter measured. The data suggests that storage and composting are optimal for the removal of human pathogens and anaerobic digestion for the reduction in AMR genes and mobile genetic elements. The nitrogen content is increased in storage and AD and reduced in composting. Thus, depending on the requirement for increased or reduced nitrogen the optimum treatment varies. Combining the results indicates that composting provides the greatest gain by reducing risk to human health and the environment. Network analysis revealed reducing Proteobacteria and Bacteroidetes while increasing Firmicutes will reduce the AMR content. KEGG analysis identified no significant change in the pathways across all treatments. This novel study provides a data driven decision tree to determine the optimal treatment for best practice to minimise pathogen, AMR and excess or increasing nutrient transfer from slurry to environment.

**Graphical abstract:** 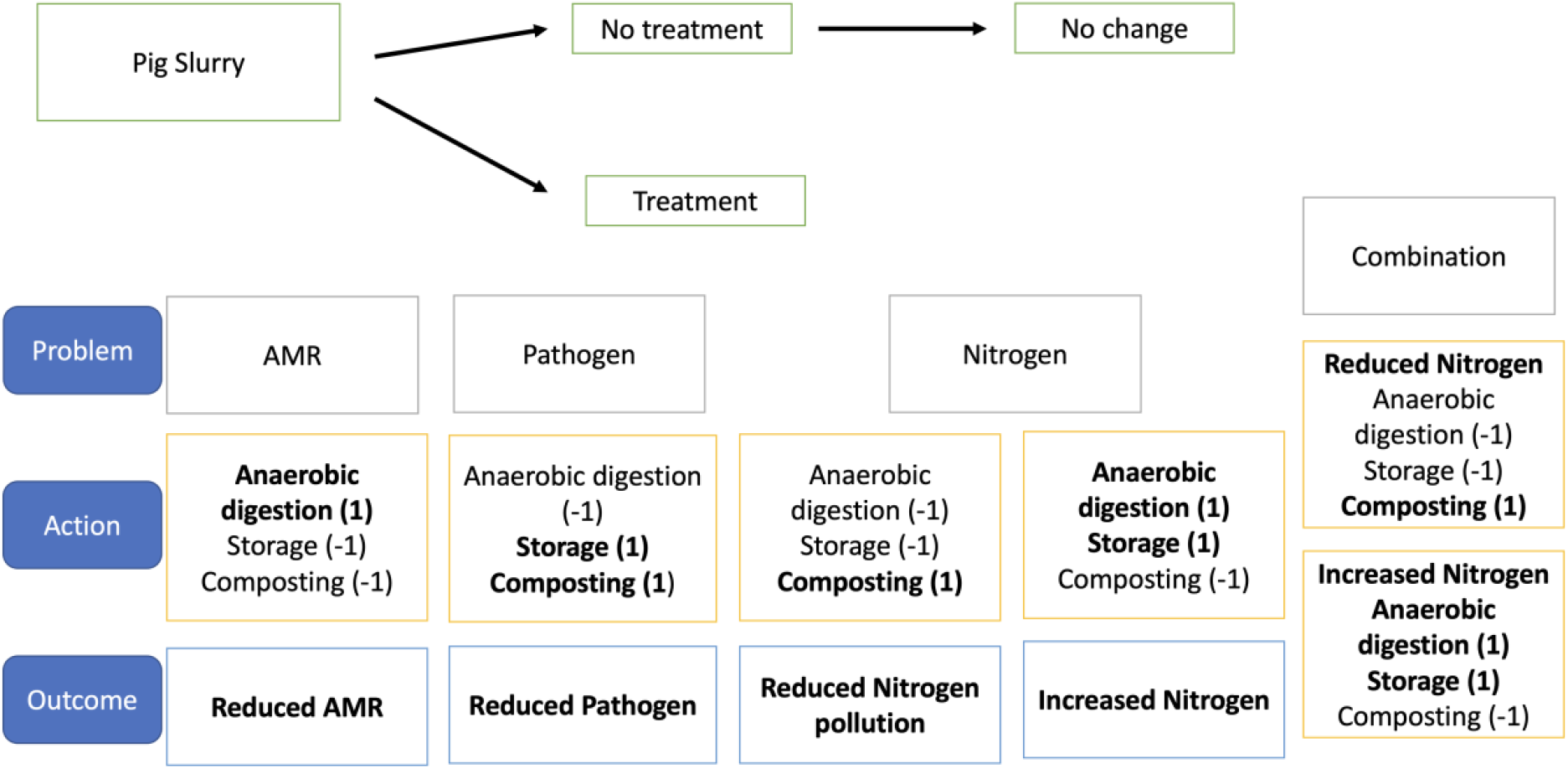

## 1. Introduction

Antibiotic use in human and veterinary medicine plays a key role in antibiotic resistance (AMR) dissemination. In 2015 the World Health Organisation released their Global Action Plan on AMR to tackle the AMR crisis in both human and animal health. Within the EU, the European Medicines Agency encourages the careful use of antibiotics in humans and animals and started a programme to collate data concerning antibiotic use. Globally, tetracyclines, penicillins and macrolides are the most commonly utilised antibiotics in pig production (Lekagul et al., 2019). The presence of antibiotics at very low concentrations (up to several hundred-fold less than the breakpoint concentrations for pathogens) can select for antibiotic resistance (Gullberg et al., 2011). The development of resistance among bacteria may occur in the gastrointestinal tract of animals, and in different environmental niches (e.g., organic fertilisers, agricultural land, different water sources). The antibiotic resistance genes (ARGs) present on mobile genetic elements (MGEs) in antibiotic resistant bacteria (ARB) can be transferred to antibiotic susceptible bacteria within a wide range of biomes (Rasschaert et al., 2020).

Pig slurry contains antibiotic residues, AMR bacterial pathogens and commensals (Rasschaert et al., 2020). The application of animal slurry as organic fertilisers has been shown to impact the soil microbial communities and be the main factor shaping the antibiotic resistome therein (Jechalke et al., 2014; Udikovic-Kolic et al., 2014). According to the One Health concept, environmental biomes connect with human and animal biomes, thus the ARB and ARGs in slurry can transmit to humans and animals and transfer to environmental niches when used as soil fertilisers. High abundances and diversities of ARGs have been detected in pig slurry (Zalewska et al., 2021).

Farmers use pig slurry as fertiliser spread on agricultural farming land to improve crop production, minimise fertiliser costs and contribute to the circular economy. However, the application of pig slurry may result in serious environmental issues such as the risk of introducing pathogens, ARB, and ARGs to the soil, crop, and increased nutrients such as nitrogen in water sources (Peu et al., 2006; Rasschaert et al., 2020; Zhu, 2000). Slurry treatment methods such as composting, anaerobic digestion (AD) and storage have been used to reduce the antibiotic residues, ARB, and ARGs in slurry before application to soil as an organic fertiliser (Li et al., 2020; Tran et al., n.d.; Zhou et al., 2019). Compost treatment of slurry is a cost-effective technique that can be implemented as an environmentally friendly method of recycling pig slurry. The high quality compost showed a clear benefit when used as a soil fertiliser based on reduction in biological risk and high nutrient contents (Rao et al., 2007). The composition of microbial communities in composting strongly depend on the initial microbiota of the compost material and the composting process (Cao et al., 2020). In one study the dynamic change in the community composition during composting pig slurry with sawdust increased with the compost fermentation. The composting procedure showed different effects on ARGs in pig slurry. Indeed, in that study the total ARGs increased during pig slurry composting (Cao et al., 2020). However, the relative abundances of individual ARGs varied, with some having an increase in their relative abundance, while others were decreased (Chen et al., 2007; Selvam et al., 2012; Wang et al., 2015; Zhang et al., 2017).

Anaerobic digestion treatment of pig slurry, results in the reduction of greenhouse gas emissions, high-quality gas production, and is environmentally friendly, leading to a decrease in water and air pollution (Massé et al., 2011). It is considered an efficient management method of pig slurry for indoor intensive pig farms. This treatment provides heat for the pig houses where the pig manure can be continuously collected all year. The AD product, digestate, has been used widely as soil fertiliser because of its low cost compared with other approaches such as liquid-solid separation, pyrolysis of the solid fraction, and wastewater treatment for the liquid fraction (Camilleri-Rumbau et al., 2021; Zhang et al., 2021). Research has shown the fate of ARGs after AD treatment was dependent on AD starting modes, and most of the genes were carried by Firmicutes (Zhi et al., 2021). Moreover, the diversity of bacteria was at a higher level than Archaea in AD. The bacterial community was dominated by Firmicutes, while Euryarchaeota was the most abundant among Archaea (Zhi et al., 2021).

The storage of pig slurry can reduce the quantity of faecal indicators present and can significantly change the microbial community composition (Peu et al., 2006). The effect of pig slurry storage on the dynamics and characteristics of the *Escherichia coli* population was determined in the work of Duriez and Topp (Duriez and Topp, 2007). Indeed, a higher diversity was observed in the *E. coli* population in the stored manure than in the fresh faeces. The microbial communities of pig manure also changed depending on the storage duration and stored temperature (Hwang et al., 2016; Lim et al., 2018). Thus, across each treatment the input and specific treatment conditions affected the composition of the final treated products.

Previous studies of microbiota and resistome changes in pig slurry were explored either within one treatment method, within modified treatment conditions or without any treatments. However, a comparative study across different treatment methods of the same initial slurry has not been previously undertaken. The comprehensive analysis of the pig slurry microbiota in different recycling methods is required to understand the dynamics of the microbiota and its resistome when pig slurry is used as soil fertilisers. Here, we present the first study globally comparing the impact of three treatments (storage, compost, and AD) of pig slurry on the microbial community dynamics and resistome. We hypothesised that different treatments applied to pig slurry would result in the substantial changes in the human pathogen content, resistome, microbial community composition, functional pathways, and nutrient content in pig slurry. We used a combination approach of -omics and molecular biology techniques (shotgun metagenomic sequencing and HT-qPCR arrays) to explore and compare the effect of storage, compost, and AD treatments on the dynamics of microbial community composition, functional and metabolic pathways, and resistance gene profiles of the same initial pig slurry. Correlation analysis was applied to identify the links within microbial communities, between the microbial taxa, functional pathways, and resistance genes in slurry samples. We aimed to identify the most applicable and effective treatment currently available to farmers to reduce AMR transmission and minimise the onward risk to the environment, animals, and humans. Based on the data generated we aimed to also provide a matrix decision tree to determine the optimum treatment depending on the most significant problem to be addressed or by addressing a combination of all problems, but with the chance of compromise increasing with each additional problem tackled.

## 2. Materials and Methods

### 2.1. Sample collection

Pig slurry was taken from a house with 14 piglet pens, each pen contained 28 pigs. At the different stages, the diets for pigs may contain zinc oxide, Stabox and pulmotil (200 ppm per ton), and CTC (15%). The pig slurry used in this study was collected from an Irish pig farm (weaning house) in April 2019 in 3 intermediate bulk containers (IBC tanks). Two tanks of samples were used for AD (in triplicates) and the third tank was use for the compost treatment.

### 2.2. Pig slurry treatments

#### 2.2.1. Storage and compost treatments of pig slurry

In the storage treatment, pig slurry was stored in 3 plastic drums (100 L) with a lid and housed in a Teagasc research facility for 4 months in an open shed subjected to ambient air conditions. The slurry drums were manually mixed every 4 days.

The IBC tank was let to settle for 3 weeks to collect the solid fraction in the bottom of the tank. To establish the composting treatment the solid fraction of pig slurry was mixed with locally sourced sitka spruce sawdust at the ratio 4 (slurry):1(sawdust) and resulted in a total wet weight of approximately 150kg in each 200 L blue plastic drum. The composting procedure was performed in triplicate on a Teagasc research facility inside a shed. The drums containing compost mixtures were covered with lagging jackets to keep the temperature during the composting process. Compost mixtures were turned after 4 days, 1 week, 2 weeks, 3 weeks, 4 weeks, 5 weeks, 6 weeks, and 8 weeks. The temperature was recorded at the depth of 30 – 40 cm from the surface of each compost drum before every turn. The temperature of the compost within the drums reached more than 50 °C from day 6 and reduced to greater than 49 °C on day 14 (Figure S1).

Samples were collected every two weeks during all treatment processes. Slurry samples in storage drums were mixed to homogenise before each sampling time. Compost samples were collected during a composting process every two weeks after turning. Collected samples were transported to the laboratory for microbiology and molecular biology studies.

#### 2.2.2. Anaerobic Digestion of pig slurry

The AD was performed in the Microbiology Department at NUI, Galway. The AD comprised triplicate 10 L laboratory-scale continuously stirred tank reactors (CSTRs) were operated at 37 °C and fed pig slurry co-digested with fats, oils, and grease (FOG), at an organic loading rate of 2 g VS L^−1^ d^−1^ and a target retention time of 28 days (Figure S2). A three-day semi-continuous feeding regime was used for the duration of the 90-day trial. The FOG was sourced from an AD plant in Co. Kilkenny, Ireland, collected in a 25 L drum, stored at 4°C and mixed thoroughly before use. To prepare feedstock, pig slurry was mixed with FOG at a 2:1 PS:FOG ratio, based on the mixing ratio used in practice at the source farm-based AD plant. The mixed feedstock was tested at each time point for total and volatile solids (TS & VS), pH, total and soluble chemical oxygen demand (tCOD & sCOD), NH_3_-N (Table S1), before being fed through the feeding port on top of each bioreactor. Volatile solids of the input feedstock were determined regularly to account for any fluctuations in solids content between PS collections. Hence, in a 21-day cycle, bioreactors received an average of 42 g VS L^−1^ (8 L working volume), regardless of feeding regime employed. The digestate produced was analysed for TS, VS, pH, NHc-N, tCOD and sCOD. Biogas produced in the 10 L bioreactors was collected in 25 L Tedlar SCV gas bags and volume was determined at each sampling point using water displacement. Methane content of the biogas was analysed using a Varian 450 gas chromatograph (GC) equipped with a flame ionisation detector. The carrier gas was nitrogen, and the flow rate was 25 mL/min. Analysis of total and volatile solids (TS/VS) was performed gravimetrically according to standard methods (APHA, 2005) and total and soluble chemical oxygen demand (tCOD/sCOD) analysis was performed according to the Standing Committee of Analysts (Standing Committee of Analysts, 1985). NH_3_ concentrations (mg/L) were determined using the HACH AmVer High-Range Ammonia test, following the manufacturer’s instructions. The concentrations of all elements decreased after AD. The results of key physicochemical parameters were recorded throughout the 90-day AD trial (Supplementary Material Figures S3 – S6). Total and volatile solids in the digestate declined over time as the digesters had previously been fed pig slurry with higher solids content than the pig slurry (Figure S3). Ammoniacal nitrogen (NH_3_-N) in the digestate remained within a range of 2700-3500 mg L^−1^ throughout the trial (Figure S4), whilst pH also remained stable throughout. Following an adjustment phase of approximately 30 days, methane (CH_4_) production stabilised and remained consistent from Day 45 to the end of the trial, with consistently good quality biogas production (Figure S5). Total chemical oxygen demand (COD) in the digestate increased over the first 30 days but stabilised in line with stabilisation of biogas production (Figure S6), with an average 77.3% tCOD removal and 76.4% sCOD removal.

### 2.3. Physiochemical compositions of pig slurry used

Dry matter (DM) of pig slurry was assessed by oven drying of a known wet weight of sample at 105 °C for 24 hours. The dry matter content was determined by the following formula:

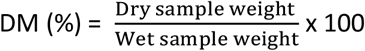

The organic matter (OM) was determined by ignition of the dry weight at 550 °C for 2 hours and measuring the weight of the ash. The OM was calculated by the expression:

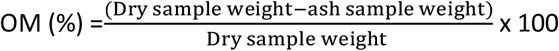

The concentrations (ppm) of available nutrients (phosphorous (P), Potassium (K), magnesium (Mg), sulphur (S), sodium (Na), and calcium (Ca)) were measured by IPC-OES using an Agilent 5100 ICP-OES spectrometer (USEPA and EPA, 1996). The samples were also freeze dried for total carbon (TC) and total nitrogen analysis by high temperature combustion method with a LECO TruSpec analyser.

### 2.4. Bacterial enumeration from compost, AD, and storage samples

The enumeration of potential human and animal pathogens (*Escherichia coli, Klebsiella* spp., *Acinetobacter* spp., *Pseudomonas* spp., *Enterococcus* spp., and *Staphylococcus* spp) was performed from pig slurry before and every two weeks during each of the treatments: storage, compost, and AD. At each sampling time six potential bacterial pathogens were enumerated from 1g of sample material from each treatment using selective agars with appropriated antibiotics (Table S2). Briefly, 1g of each sample/sample pellet was added to a 50 mL falcon tube with 10 mL of phosphate buffered saline (Gibco) and homogenised by shaking for 30 minutes. The appropriate decimal dilutions of samples were plated on selective agars with and without antibiotics, followed by incubation at 37 °C overnight up to 48h. The selected bacteria were identified based on their phenotype on selective agars according to the manufacturer selection guidelines for each agar type.

### 2.5. DNA extraction

The total DNA was extracted from 0.25g of each sample/sample pellet using the DNeasy PowerSoil Kit (Qiagen). The quality and quantity of extracted DNA were examined using a DeNovix DS-11 spectrophotometer and an Invitrogen Qubit Fluorometer (dsDNA high-sensitivity assay kit) (Waltham, MA). DNA was extracted in triplicate from each sample and extracts were pooled to obtain a single DNA sample per experimental unit at each time point. A total of 20 pooled DNA samples were subjected to metagenomic sequencing (Liverpool) and HT-qPCR arrays.

### 2.6. Metagenomic sequencing and HT-qPCR arrays

The pooled DNA samples were used to prepare a shotgun sequencing library with Illumina the NEBNext Ultra II Kit (~350 bp inserts) before sequencing on the Illumina NovaSeq 6000 platform using SP chemistry (paired-end, 2×150 bp sequencing) in the centre for genomic research, Liverpool.

The HT-qPCR arrays were performed using the SmartChip™ Real-Time PCR system (TakaraBio, CA, USA) by Resistomap Oy (Helsinki, Finland). The mix of DNA samples with primer sets and the qPCR reagents were loaded in each 100 uL reaction well of the SmartChip™ with 5182 wells. A primer set of 216 pairs of primers targeted 186 ARGs conferring resistance to major antibiotic classes, 6 integrons, 22 MGEs, and total bacterial genes 16S rRNA was used in the qPCR array. The melting curves and Ct values were analysed using default parameters of the SmartChip™ qPCR software. The qPCR was conducted in three technical replicates for each DNA sample.

### 2.7. Data analyses

The pig slurry samples were classified into 4 groups for metagenomic analysis, which included (1) a control sample group (F0-1, F0-2. F0-3, F0-4, and F0-5) collected prior to treatments: contained raw pig slurry samples collected before any treatment. There were 4 control samples F0-1 to F0-4 used in the HT-qPCR array; (2) a storage sample group (PS-W2 to PS-W16): samples collected every 2 weeks during 16 weeks of the storage treatment; (3) an anaerobic digestion (AD) sample group (AD-W1 to AD-W16): samples collected at week 1, week 2 and every 2 weeks after that until week 16 of AD process; (4) a compost sample group (CP-W2 to CP-W8): samples collected during pig slurry composting at weeks 2, 4, 6 and 8.

#### 2.7.1. Metagenomic

The adapter sequences were trimmed from shotgun sequencing raw reads in the fastq format using Cutadaprt (V2.10). We used Sickle (v1.33) with the minimum window of quality score of 20 to remove the low-quality reads with the length less than 20bp. The FastQC (v 0.11.9) was used to examine the quality of filtered reads before assembly with Megahit (v1.2.6, --kmin-1pass –presets meta-large) (Li et al., 2015). The assembled contigs were subjected to Kaiju taxonomic classifier (v1.2.6, parameters: --kmin-1pass –presets meta-large)(Menzel et al., 2016). to assign the taxonomy profile for each sample. The microbial communities were analysed in the MicrobiomeAnalyst online platform (Dhariwal et al., 2017).

The microbial genome annotation on filtered reads was carried out using Prokka (v 1.14.6, default settings)(Seemann, 2014). The protein FASTA files resulted from the Prokka software were used to identify the KEGG Orthologs (KOs) by Kofamscan (v 1.3.0). The KEGG pathways were assigned by MinPath software (v 1.5) based on the Ko’s lists(Ye and Doak, 2009). Data were analysed and visualised using Calypso online (Zakrzewski et al., 2017).

#### 2.7.2. qPCR

The qPCR data of the samples were retained for further analysis if the data in the samples met the following criteria: (1) a gene was detected in at least two technical replicates; (2) the Ct values ≤27; and (3) the amplification efficiency was in the range of (1.8-2.2). The relative gene copy number was calculated in equation 1 of the work of Chen et al. 2016 (Chen et al., 2016). Then the gene copy number was identified by dividing the relative copy numbers by the 16S rRNA gene copy number. The data was then analysed in the Calypso online platform.

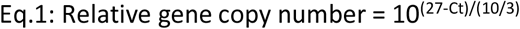

#### 2.7.3. Correlation analysis

The interaction between (1) microbial communities and ARGs, (2) ARGs and MGEs, (3) between microbial communities and KEGG functional pathways were analysed through Spearman’s correlation analysis with the SciPy package. The correlation was considered strong and significant when Spearman’s rank value |r| > 0.7 and p < 0.05. The Cytoscape software (v3.8.2) was implemented to build the network based on strong and significant Spearman’s correlations.

### 2.8. Data and material availability

The sequencing data of the swine slurry samples have been submitted in the NCBI Sequencing Read Archive (SRA) under the bio-project PRJNA771392. Relative abundances of microbial taxa, ARGs, MGEs; and correlation analysis data as well as HT-qPCR array data are available as supplementary material (Data S1-S5.)

## 3. Results

### 3.1. Microbial analysis of pig slurry samples during different treatments

*E. coli, Klebsiella* spp., *Acinetobacter* spp., *Pseudomonas* spp., *Enterococcus* spp., and *Staphylococcus* spp. were detected in the pig slurry prior to treatment and a reduction in abundance was observed in all treatments over the experimental period, to varying extents (Figure 1). *Enterococcus* spp. were the only bacterial species to persist to the final timepoints during storage and compost treatment. *Acinetobacter* spp., *Staphylococcus* spp., and *Klebsiella* spp. were detected in storage samples collected until week 12, while *E. coli* was detected until week 10, and *Pseudomonas* spp. was detected in all samples until week 8 (Figure 1a). Among compost samples, *Klebsiella* spp., *Acinetobacter* spp., *Pseudomonas* spp., and *Staphylococcus* spp. were all absent in the final week 6 sample. *E. coli* was detected in the week 2 sample only (Figure 1c). The AD treatment led to a rapid (week 1), substantial initial reduction of all analysed bacterial species in pig slurry, compared to the control samples. However, unlike in the storage and compost treatments, the six bacterial pathogens were present in all samples collected during AD treatment, including the final timepoint at week 16, meaning that the AD treatment had the highest putative pathogen load of all treatments post treatment. There was little reduction in the cfu/g of each pathogen in the AD samples from week 1 to week 16 (Figure 1b). At week 16 there was a minimum of 10^3^ cfu/g of each pathogenic species detected in the samples. Few bacterial colonies were detected on selective agars that were supplemented with antibiotics, indicating a low level of AMR in the species tested present in the slurry initially and throughout the treatments.

**Figure 1.**
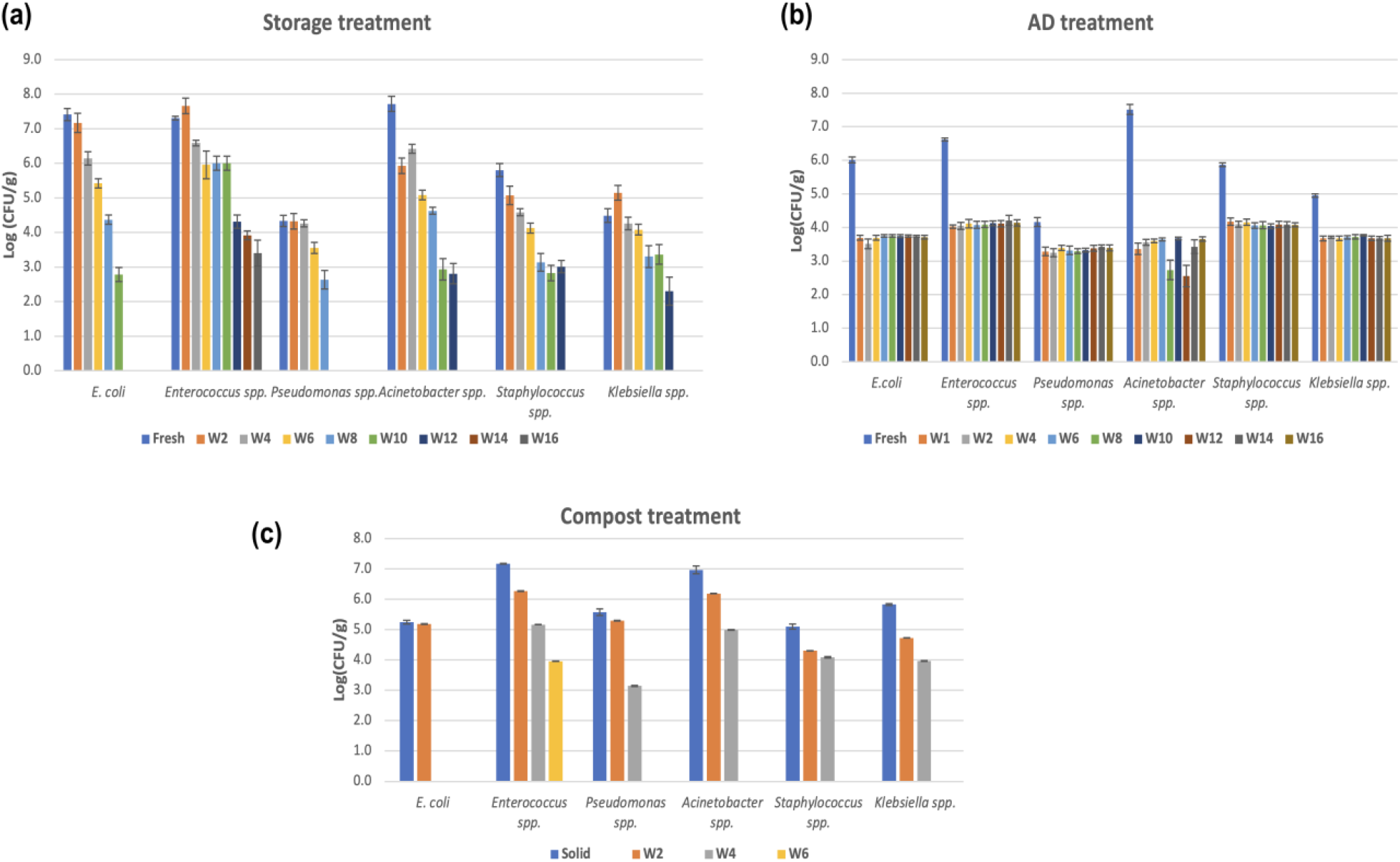
Bacterial enumeration during treatment by storage (a), AD (b), and compost (c) treatments of pig slurry. The bacteria were enumerated on agar without selective antibiotics. Fresh: raw slurry before treatments; Solid: solid fraction of pig slurry, which was used for composting with sawdust; W2-W16: weeks in which the samples were collected e.g. W2 = week 2.

### 3.2. Microbial composition changes due to treatments

After quality trimming and assembling, 40.34 to 78.44% of the metagenomic sequencing reads could be assigned to Bacteria, Archaea, Virus and other unclassified organisms with Kaiju (Table S3). The composition of microbial communities was assessed at the phylum level for all samples (Figure 2). The most abundant phylum differed across the sample groups. However, the three phyla Firmicutes, Bacteroidetes, and Proteobacteria, were dominant in all samples. The phylum Firmicutes was the most abundant phylum among control (30.67% - 55.75%), and storage (30.62% - 38.8%) samples, followed by Bacteroidetes (from 19.72% to 53.56% across control and storage samples), Proteobacteria (5.07% to 67.25% across control and storage samples), and Actinobacteria (2.2% - 8.3% across control and storage samples) (Data S1). In the AD samples, Firmicutes was also the most abundant phylum (49.99% - 60.34%), followed by Bacteroidetes (11.4% - 20.56%), Proteobacteria (6,26% - 8.54%), and Euryarchaeota (3.55% - 9.56%). While the most abundant phylum in the compost samples was Proteobacteria (48.5% - 67.2%), followed by Bacteroidetes (10.24% - 41.2%), Firmicutes (5.26% - 18.98%), and Actinobacteria (1.27% - 3.49%). The storage process showed a different effect on the relative abundances of microbial phyla (Data S1). Some phyla were decreased in relative abundances such as Firmicutes, Spirochaetes, and Euryarchaeota, while other phyla showed an increase (Bacteroides, Proteobacteria) or were consistent throughout (Actinobacteria, Spirochaetes). The decrease in relative abundances was also found for Bacteroidetes and Actinobacteria during AD, while Firmicutes and Euryarchaeota were increased. The compost treatment also had a differential effect on the relative abundances of microbial phyla, with a decrease for Firmicutes and Actinobacteria and an increase for Proteobacteria in week 2. Bacteroidetes decreased in week 2 samples but increased toward week 8, with corresponding reductions in Firmicutes and Proteobacteria. Thus, composting had the greatest impact and storage the least impact on the major phyla of the slurry.

**Figure 2.**
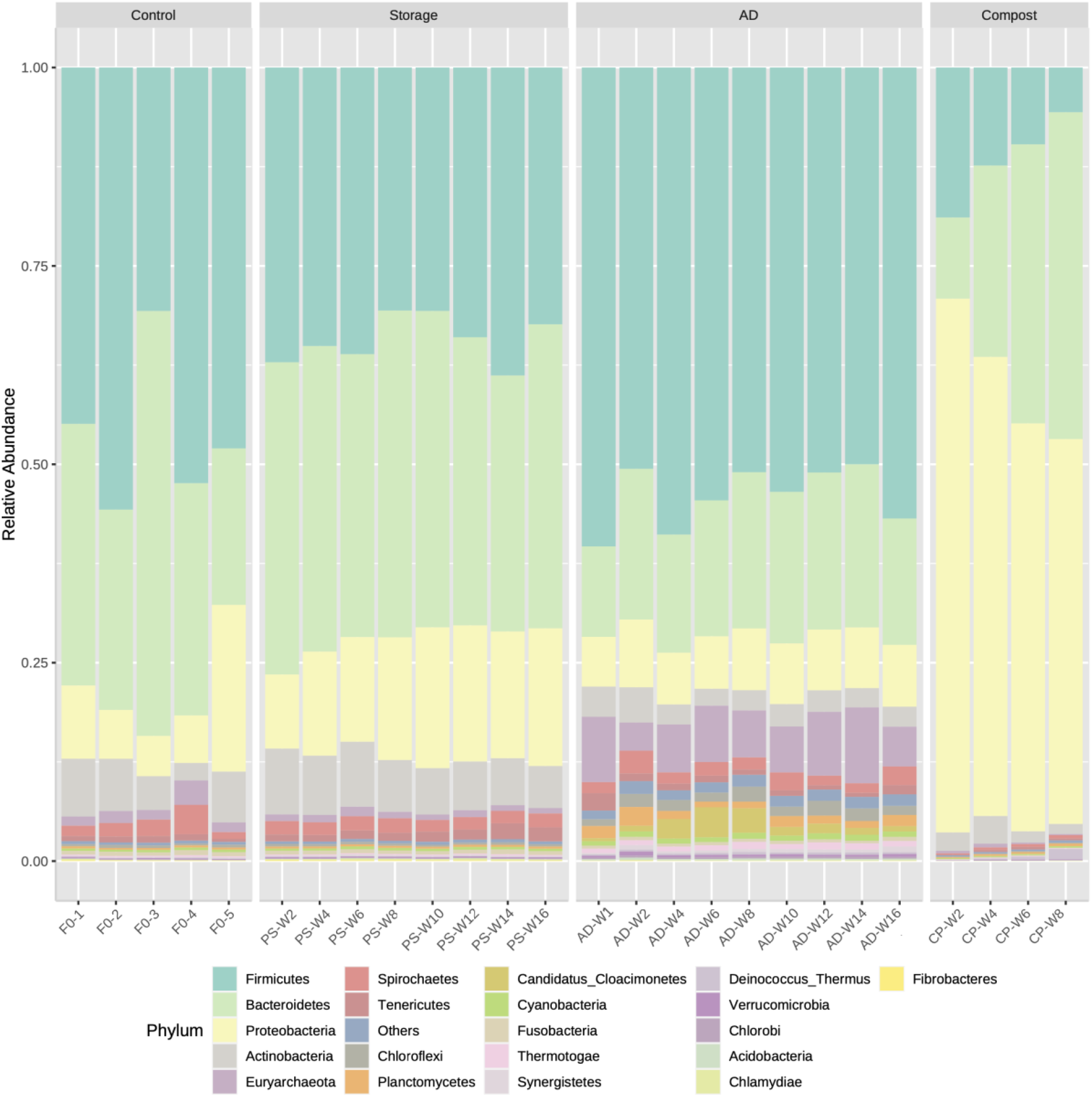
Microbial composition at phylum level. The samples were divided into 4 groups, (1) a control sample group (F0-1, F0-2. F0-3, F0-4, and F0-5) contained raw pig slurry samples collected before any treatment; (2) a storage sample group (PS-W2 to PS-W16): samples collected during the storage treatment at weeks 2, 4, 6, 8, 10, 12, 14, and 16; (3) an anaerobic digestion (AD) sample group (AD-W1 to AD-W16): samples collected every 2 weeks during 16 weeks of AD process; (4) a compost sample group (CP-W2 to CP-W8): samples collected during pig slurry composting at weeks 2, 4, 6, and 8.

The heat map of the genus profile, based on their relative abundance, also revealed the prevalence of a distinct pattern based on the treatment groups (Figure 3a). In control and storage treatments, *Prevotella* and *Bacteroides* were the most prevalent. *Hungateiclostridium* and *Syntrophomonas* were dominant among AD samples, while *Pseudomonas* was dominant in compost samples. A linear discriminant analysis (LDA) effect size (LEfSE) was performed to characterise the microbiota of pig slurry in different sample groups. This method supports high-dimensional class comparisons in metagenomic analysis. The LDA model predicts the features (microbial taxa, functional pathways) most likely different between treatment samples groups, based on their abundances and estimates the effect size of significant different taxa/pathways (Segata et al., 2011). The microbial taxa at the genus level with LDA score [log 10] > 2 among 50 of the most prevalent taxa were calculated (Figure 3b). LEfSE identified 49 representative genera, which displayed statistically significant differences of microbiota between different sample groups. The control group was characterised by 11 genera with the most enriched genera being *Prevotella*, *Lactobacillus,* and *Clostridium*. Among 12 representative genera in the storage sample group, *Bacteroidetes*, *Parabacteroidetes*, and *Corynebacterium* were the most abundant. *Syntrophomonas*, *Hungateiclostridium*, and *Methanobacterium* were the most enriched among 15 genera designated to AD samples, while the compost samples indicated 10 differential genera with the most abundance of *Pseudomonas*, *Alcaligenes*, and *Brevundimonas*. These data identify the specific genera changes due only to the treatments applied and the resulting fingerprint of the treated slurry samples.

**Figure 3.**
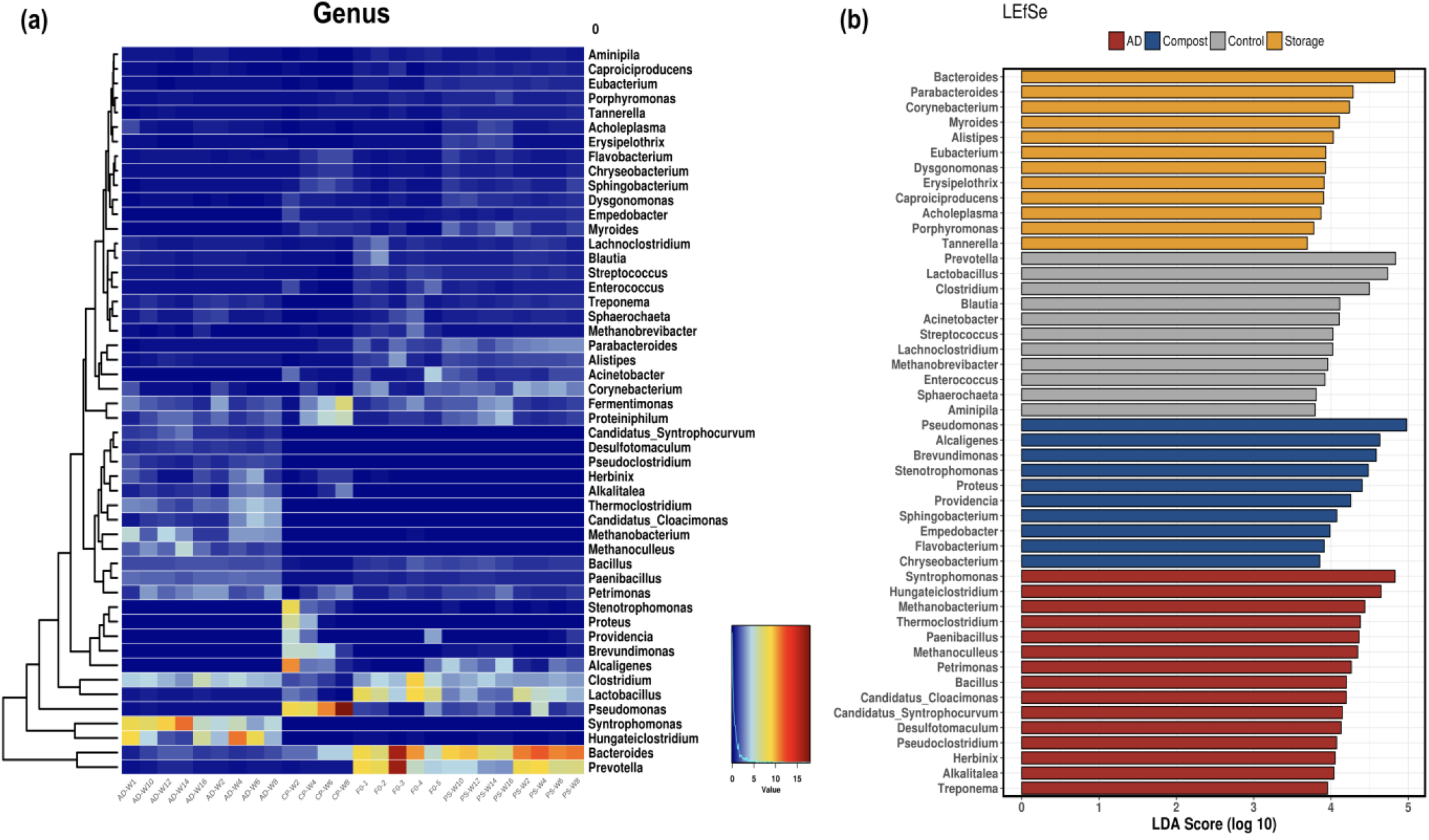
Heatmap showing the relative abundance of microbial genera (a) and characteristics of pig slurry microbial communities across control, storage, AD, and compost samples with LEfSE (b), microbial taxa with LDA score >2 among 50 the most abundant taxa are displayed.

### 3.3. Microbial community diversity

The microbial diversity of all samples was analysed based on relative abundances of identified taxa with detailed taxonomic paths. In total 7776 taxa were assigned across all samples. The richness and diversity of the microbial communities were assessed using Chao 1 and Shannon indexes (Figure 4). The Chao 1 indexes showed significantly higher richness of the microbial community in the pig slurry of the control and storage sample groups, in comparison with AD and compost samples (p <0.05). After AD and compost treatment, the Shannon diversity indexes decreased, while this value was higher in the storage sample group than those in the control. The microbial communities from all samples were also visualised by principal coordinate analysis (PCoA) based on the Bray-Curtis dissimilarity matrices (Figure S7a). The PCoA plot identified 4 differential clusters formed by 4 sample groups with an overlap only between 2 clusters of control and storage samples. These data again demonstrate the unique microbial populations within the AD and compost treated samples and the similarity between the stored and control slurry. Regardless of the time of treatment the samples clustered closer to the treatment specific samples than the control or storage, thus indicating a rapid change in the microbial composition due to treatment, which is maintained over time.

**Figure 4.**
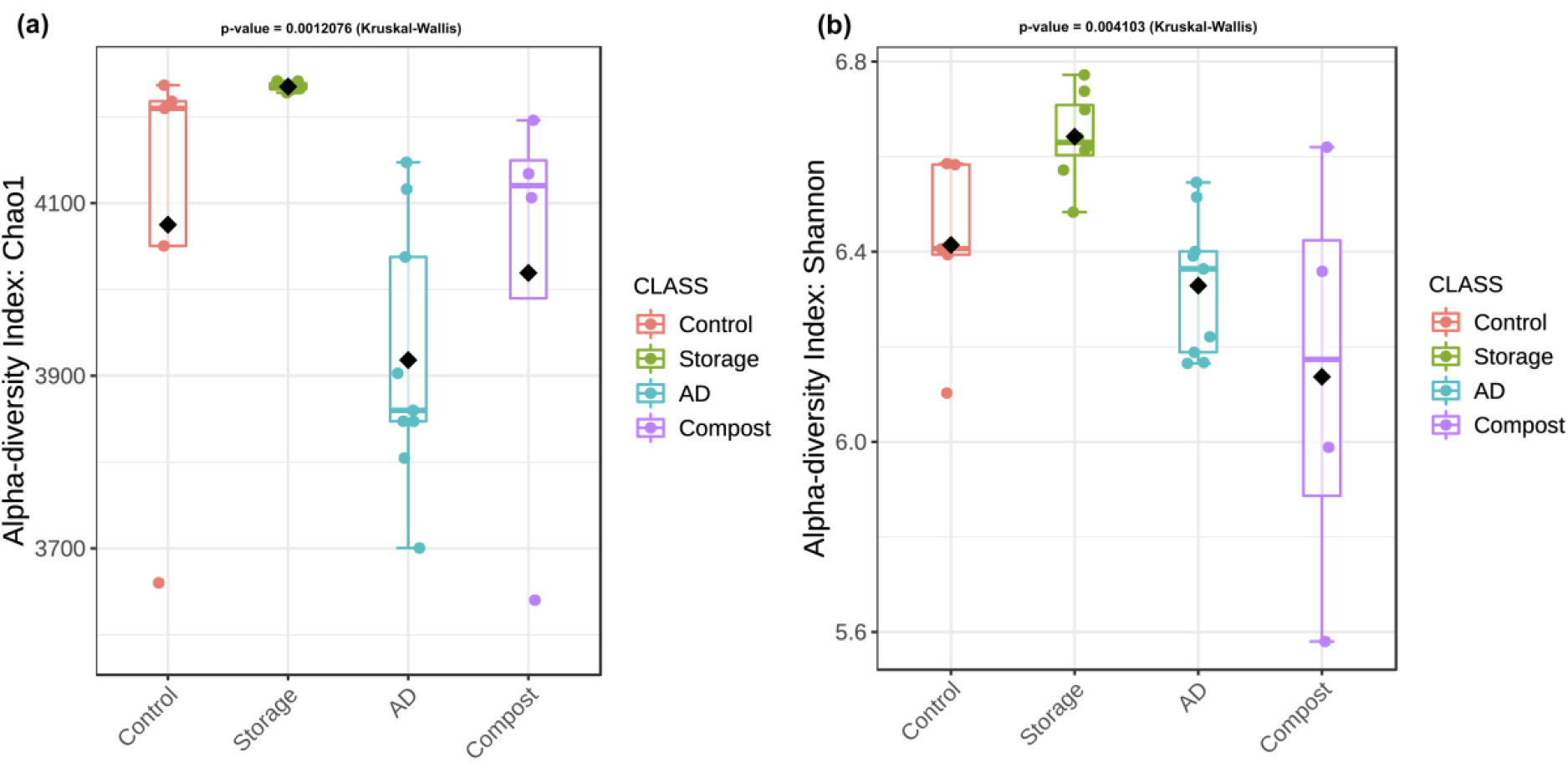
Alpha diversity with Chao 1 (a) and Shannon (b) indexes of microbial communities during control, storage, AD, and compost treatments.

### 3.4. Characteristics of the resistome in pig slurry under different treatments

A total of 181 genes comprising 154 ARGs and 27 MGEs were detected across all samples. The detected ARGs were divided into 11 main resistance groups: aminoglycoside, beta-lactam, MDR, macrolide-lincosamide-streptogramin B (MLSB), polymyxin, phenicol, quinolone, sulfonamides, tetracycline, trimethoprim, and vancomycin. Four MGE groups were identified consisting of integrons, transposons, insertional sequences, and plasmid-associated genes. The total detected genes (both ARGs and MGEs) in pig slurry increased during storage and compost treatments (Figure 5). In control sample groups, the most abundant ARGs were tetracycline, aminoglycosides, sulfonamides, and MLSB (Data S2). These ARG classes were also dominant among storage samples. The relative abundance of the total detected genes in pig slurry decreased during AD treatment. In comparison with the control samples, the final product of the AD treatment showed a decrease in the relative abundance of analysed resistance gene classes (except polymyxin resistance genes, which showed an increase from 0.0004 to 0.0008 and trimethoprim resistance genes remained at the same level of abundance). The greatest abundances were identified across aminoglycoside, tetracycline, vancomycin, and beta-lactam ARGs in the AD sample group. In the composting process of pig slurry, most of the gene classes (sulfonamides, aminoglycosides, trimethoprim, MLSB) were increased at week 2 and week 4, then reduced toward the end of the process, but remained relatively high. The most prevalent gene classes in all compost stages were resistance to sulfonamides, followed by aminoglycoside, and tetracycline.

**Figure 5.**
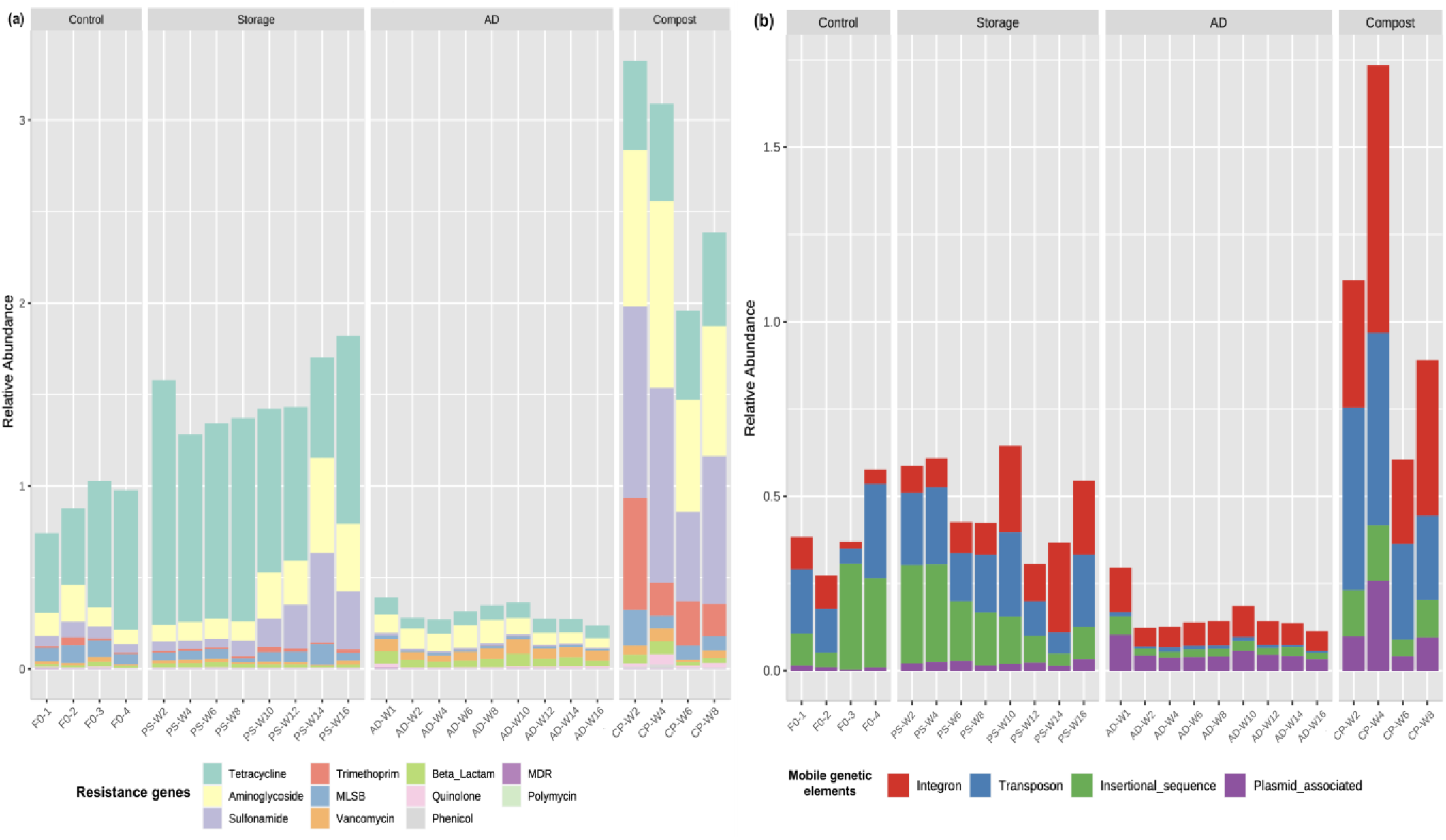
Gene relative abundance detected in pig slurry under different treatments for (a) resistance gene and (b) mobile genetic elements. The data is presented as the sum of relative abundances ARGs or MGEs. The samples were divided into 4 groups: control contained slurry samples collected before treatment (F0-1, F0-2, F0-3, F0-4); Storage (PS-W2/W16) were samples collected every 2 weeks during storage; AD were samples collected every 2 weeks during anaerobic digestion; and Compost were samples collected during composting process at weeks 2, 4, 6, and 8.

The relative abundance of detected MGEs varied in all samples. The MGEs were found at a different level across control samples. However, the ranking of the different MGE types remained unchanged. The most prevalent MGE was insertional sequences, followed by transposons, integrons, and plasmid-associated genes. A decrease of total detected MGEs were found at weeks 6, 8, 12, and 14 among storage samples. The relative abundance of insertion sequences decreased after 16 weeks of the storage treatment, while the increase was identified for other MGE classes. The dynamics of MGEs within bacterial populations is not well described. These data suggest that, with a reduction in insertion sequences, a concurrent increase in transposon and integrons occurs at week 16. The MGE trend in storage samples does not mirror the ARG data, suggesting that the MGEs contribute only partially to the resistome and ARGs could also be carried on non-mobile elements in these samples. AD led to a decrease in total MGE relative abundance in pig slurry compared with the storage sample and with the control sample groups. Among AD samples, the most prevalent MGE class was integrons, followed by plasmid-associated genes, insertional sequences, and transposons.

In contrast to AD treatment, composting of pig slurry resulted in an increase in total MGE relative abundance compared with other sample groups. The MGEs’ relative abundance increased from week 2 to week 4, then decreased in the later timepoints, but remained relatively high. The composition of MGEs in compost samples was dominant by integrons, followed by transposons, insertional sequences, and plasmid-associated genes (Data S2). The resistome data of AD and compost samples (weeks 6 and 8) mirror the MGE data suggesting an important role for the MGEs in the resistome of the AD and composting samples. This contrasts with the storage samples where more fluctuation in the MGE data occurred in comparison with the resistome data.

The richness (Chao1) of ARGs decreased in AD and storage samples (Figure 6a). However, the differences in ARGs richness were not statistically significant (p > 0.05). The Shannon indexes showed significant distinction among sample groups (p<0.05), with an increase of ARG diversity in compost and AD samples and a decrease in storage samples in comparison with the control group (Figure 6b). A significant reduction in the MGE richness was observed in all treatment groups (Figure 6c) (p<0.05). The MGE diversity was also significantly different between all group samples (Figure 6d) (p<0.05).

**Figure 6.**
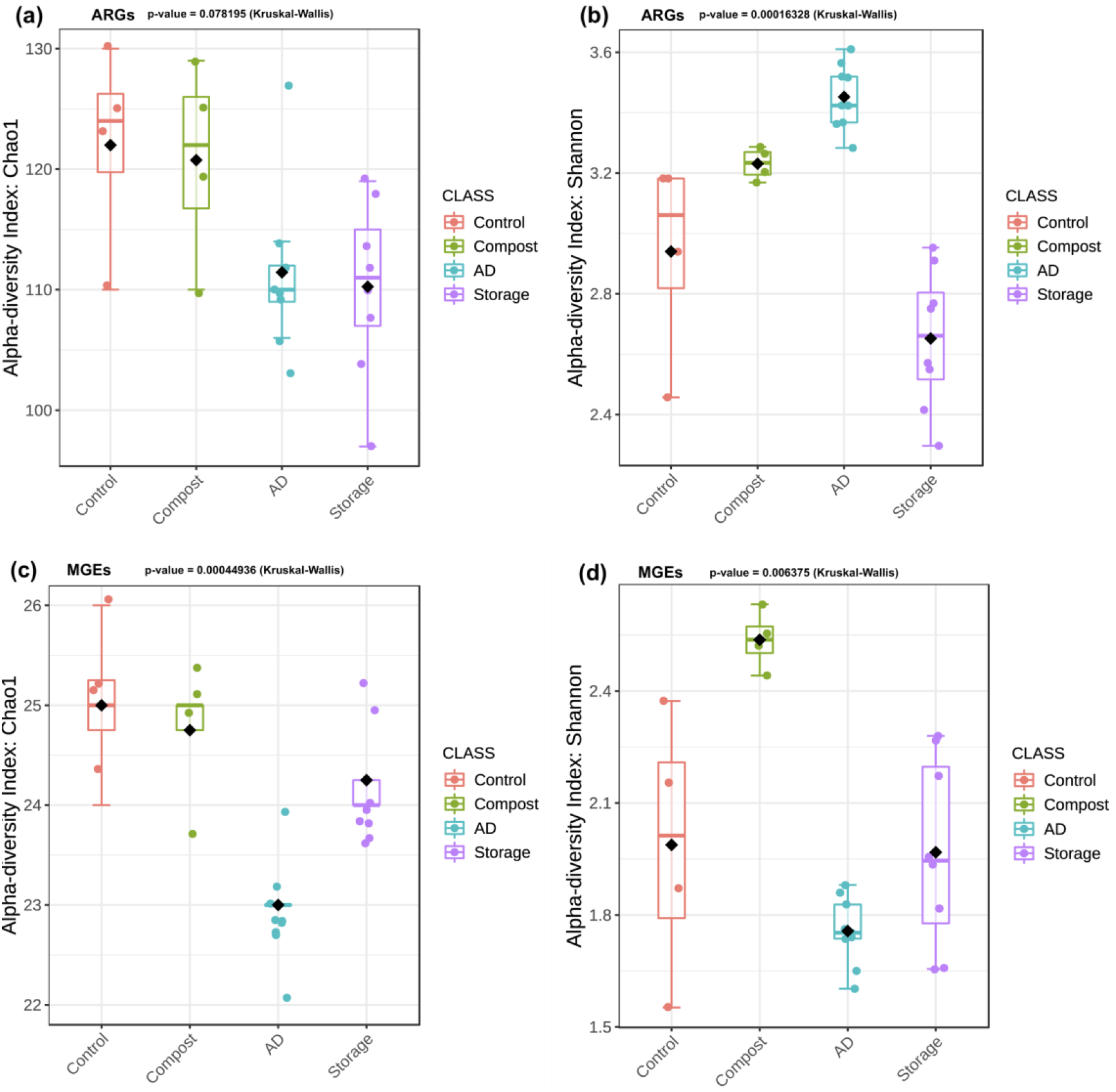
Alpha-diversity indexes: Chao 1 (richness index) and Shannon diversity indexes of ARGs (a), (b) and MGEs (c), (d) in control and treated sample groups.

The composition of ARGs and MGEs was analysed through principal coordinate analysis (PCoA) based on Bray-Curtis dissimilarity (Figure S7b, c, PERMANOVA test, p <0.05). In the ARG PCoA, control, compost, and AD clusters separated from each other, but the storage cluster overlapped with control and compost clusters (Figure S7b). The MGEs of the AD sample group also formed a cluster separated from other sample groups. The MGEs of the control, storage and composted samples overlapped with each other (Figure S7c). A core resistome containing 89 genes was identified based on the presence of these genes across all sample groups (Figure S7d). These genes included resistance to tetracycline (n=17), aminoglycosides (n=16), beta-lactam, (including *bla*_NDM_) (n=12), sulfonamides (n=5), trimethoprim (n=2), MLSB (n=4), vancomycin (n=6), phenicol (n=1), MDR (n=2), polymyxin (*mcr1*) (n=1), quinolone (n=1) and MGEs (n=22) (Table S4). This lists the genes which were not removed by any of the slurry treatments and thus persisted regardless of treatment to the final product. The list of ARGs include *bla*_NDM_ and *mcr1*, both mobile ARGs to the last line of defence antibiotics; carbapenems and colistin, respectively.

### 3.5. KEGG functional annotation

In total 354 KEGG pathways were identified across all samples. In the control and storage sample groups, the largest groups of annotated genes were assigned to ABC transporters, followed by two-component systems and methane metabolism. The methane metabolism was dominated in the AD sample group, followed by ABC transporters and ribosome. Among the detected pathways in compost samples, the most abundant was two component systems, followed by ABC transporters and purine metabolism (Figure 7a). The richness of detected pathways was higher in all treatment sample groups compared with the control group (Figure S8a). A statistically significant difference in Shannon indexes was found between all sample groups (Figure S8b). A total of 175 core pathways were identified across all samples (Figure S8c). The control samples were the only ones not to contain unique KEGG pathways. However, the numbers of pathways unique to each treatment was between 4 and 15. All samples formed 4 separate clusters, with the control sample cluster overlapping the other three (Figure S8d). Overall, the KEGG pathway distributions across sample types did not varying greatly and the overall pattern of KEGG pathways was maintained across treatments and time. Thus, while the microbial populations and resistomes varied the major KEGG pathway genes remained stable.

**Figure 7.**
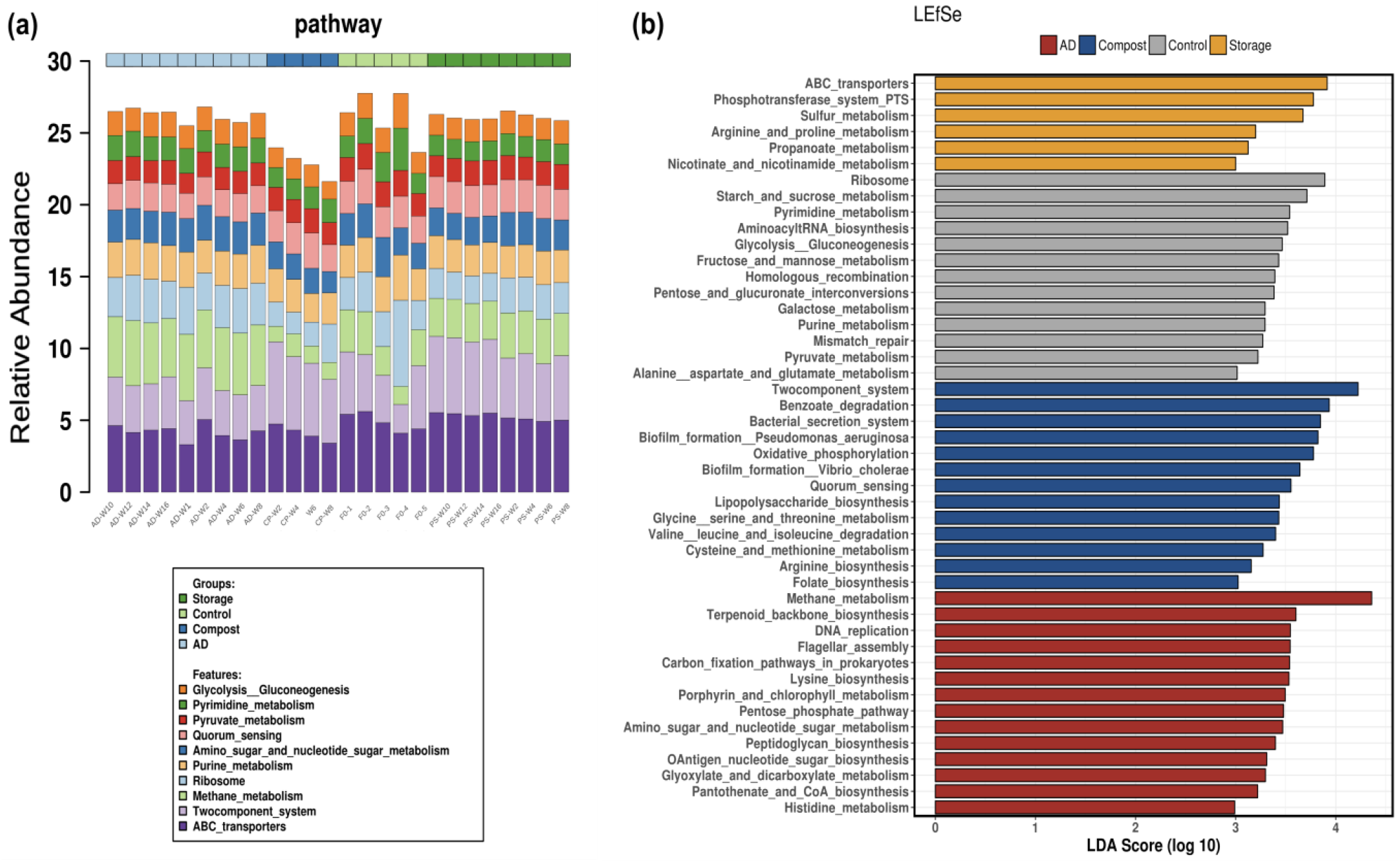
Relative abundance of KEGG pathways present in control, storage, AD, and compost samples during treatment (a) and representative pathways for each sample group with LEfSE (b), pathways with LDA score >2 among 50 the most abundant taxa are displayed.

The LEfSE analysis among 50 of the most abundant pathways revealed the characteristics of each sample group (Figure 7b). The control group was characterised by 13 pathways. Among them, the most enriched pathways were ribosome, starch and sucrose metabolism, and pyrimidine metabolism. The storage sample group was represented by 6 pathways with the most enrichment of ABC transporters, phosphotransferase system PTS, and sulfur metabolism. The compost sample group was recognised by the overrepresentation of two component systems, benzoate degradation, and bacterial secretion system over 13 representatives. Methane metabolism, terpenoid backbone biosynthesis, and DNA replication were the most enriched among 14 marker pathways of the AD sample group.

### 3.6. Network analysis

#### Association between ARGs and MGEs

The network analysis based on strong and significant Spearman’s correlations (r > 0.85, p<0.05) between ARGs and MGEs was employed to understand the co-occurrence of ARGs and MGEs across all sample groups (Figure 8, Data S3). The network was created based on 126 nodes and 489 edges formed by 93 negative (blue edges) and 395 positive (red edges) correlations. The MGEs showed a higher degree of interactions with ARGs, indicating their central role in the network formulation. The network showed the presence of two large clusters C1 and C2. In cluster 1 (C1), the *repA* and Tn5 genes displayed the most interaction with ARGs, followed by Tp614 and IS631. They formed the main hubs in this cluster. While *repA* displayed mostly positive interactions (red lines), Tn5 displayed mostly negative interactions (blue lines) with the ARGs. In the C1 cluster the interaction mainly formed between beta-lactamase resistance genes and some others including tetracycline, aminoglycoside resistance, *sul* and, *mcr-1* genes and MGEs. The main hubs in cluster C2 were formed by *tnpA* and *intI* genes (Figure 8). The C2 cluster included the interaction between MGEs mainly with aminoglycoside, *sul*, and tetracycline resistance genes. These data also demonstrate the specific ARG – MGE linkages. If all ARGs were present on all MGEs detected, the networks would have clustered based on each individual MGEs. However, these data demonstrate the varying ARG combinations associated with the same MGEs.

**Figure 8.**
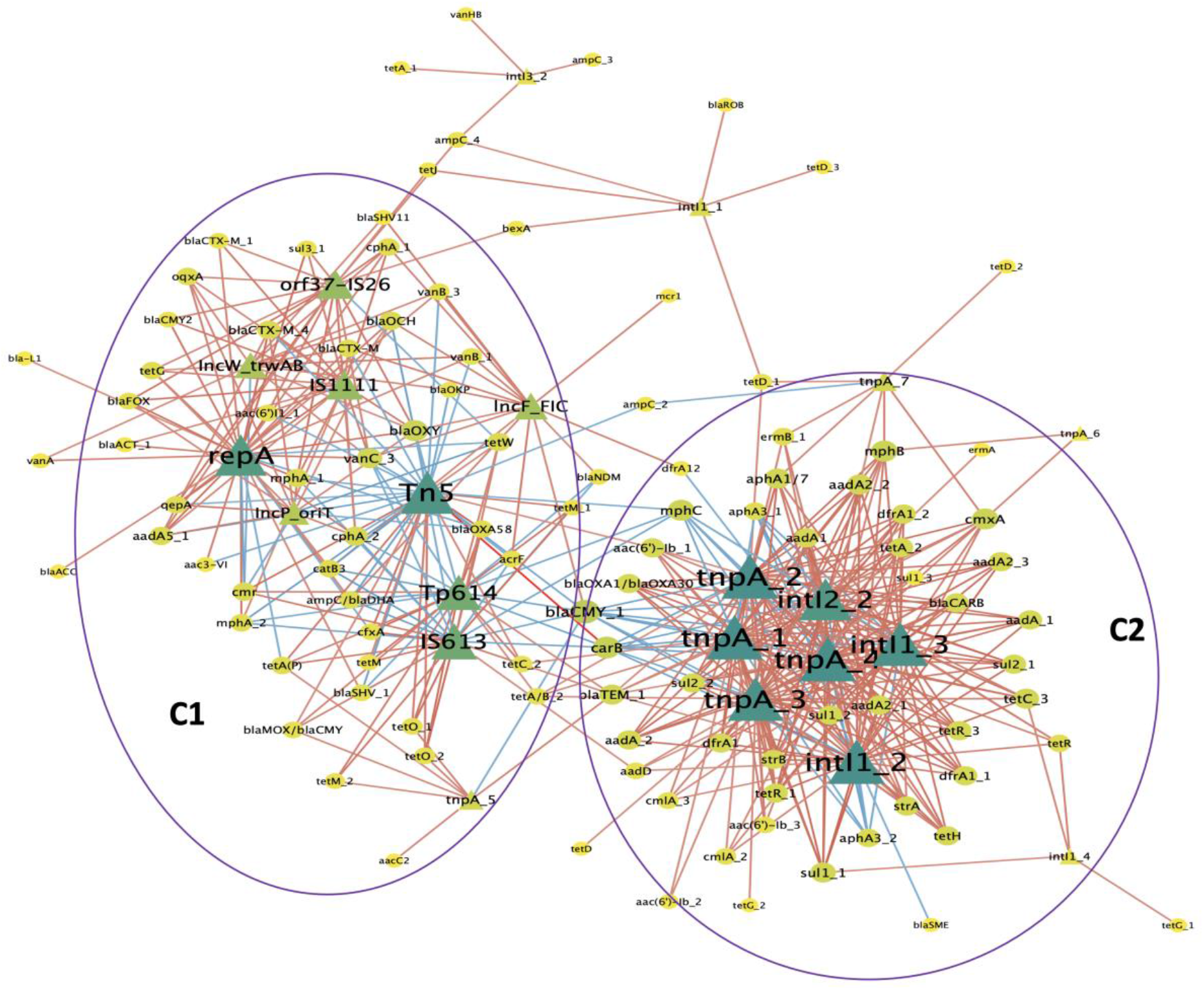
ARG and MGE interaction network analysis presented in “organic layout”. A connection shows a strong and significant correlation based on Spearman’s rank analysis (|r| > 0.85, p<0.05). The red and blue edges indicated the indexes of positive and negative correlations, respectively, between ARGs and MGEs. The size and colour (ranging from yellow to dark green) of the nodes showed degree of the interactions.

#### Correlation analysis between microbial taxa vs resistance genes and between microbial taxa vs functional pathways

The relationship and interaction between microbial phyla and ARGs were investigated in the network based on Spearman’s correlation analysis (Spearman’s |r| > 0.7, p <0.05). The network displayed strong correlations between phyla and ARGs (Figure 6b) with 61 nodes (from 7 microbial phyla and 54 ARGs), and 163 edges (built from 113 negative (blue lines) and 50 positive (red lines) correlations). Proteobacteria had the most positive interactions with 20 ARGs, followed by Bacteroidetes having positive interactions with 7 ARGs of the bacteria. Actinobacteria had positive correlations with *tetW* and *bla* _CTX-M-6_ only (Figure 9a, Data S4). These results indicated their role as primary ARG hosts. When these data are compared with the relative abundance data of phyla and ARGs across samples (Figures 2 and 5a) we can identify the trend of increasing Proteobacteria and Bacteroidetes in compost samples and increasing relative abundances of ARGs. Whereas increases in Firmicutes in the AD samples did not result in increases in ARGs. Firmicutes displayed negative correlations with 22 ARGs. Classes of genes conferring resistance to aminoglycosides (*aphA* and *aadA*) and tetracyclines (*tet*) had strong interactions with different microbial phyla (45 and 32 connected edges, respectively).

**Figure 9.**
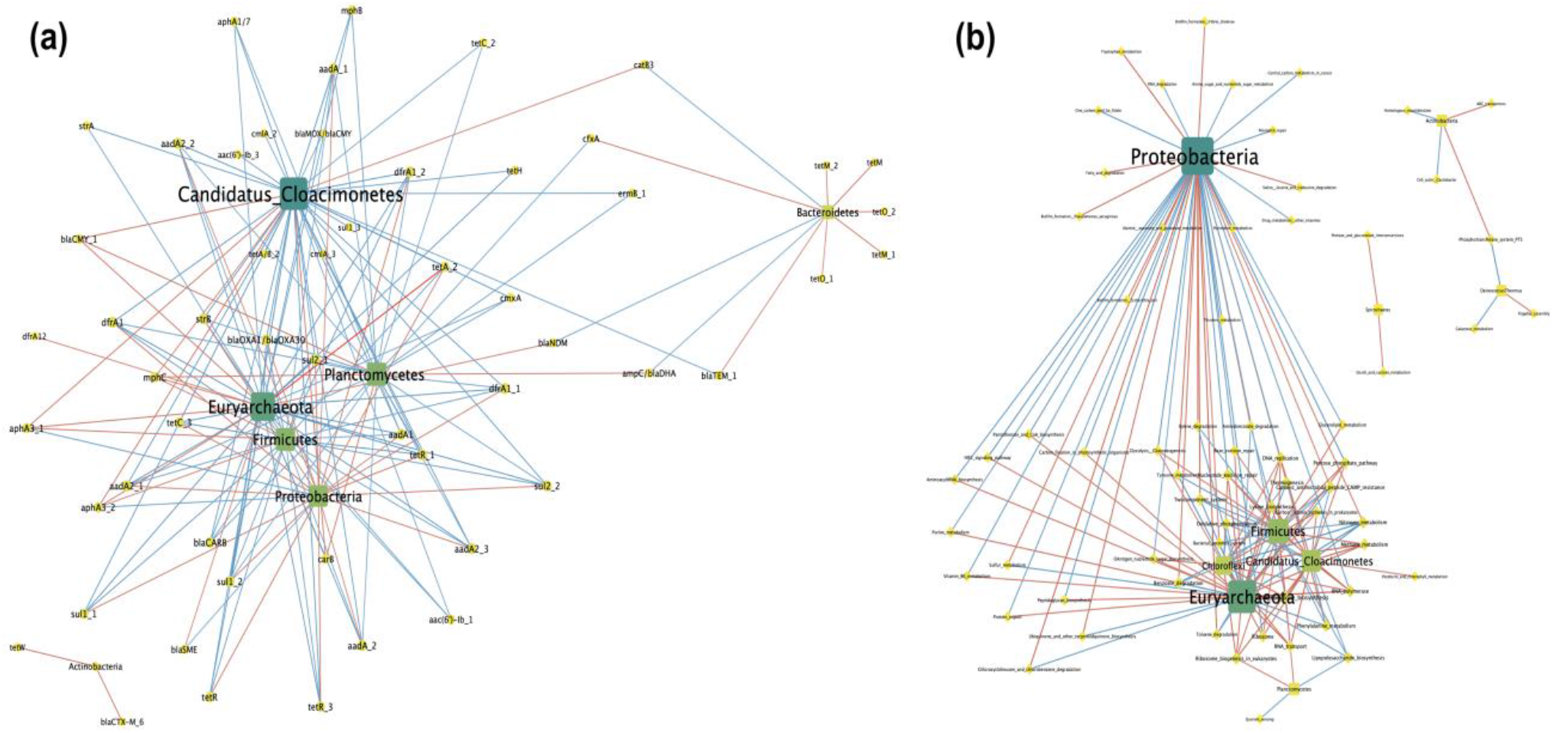
ARG and microbial phylum interaction network analysis presented in “organic layout” (a) and between microbial phyla and KEGG functional pathways (b). A connection shows a strong and significant correlation based on Spearman’s rank analysis (|r| > 0.85, p<0.05). The red and blue edges indicated the indexes of positive and negative correlations. The size and colour (ranging from yellow to dark green) of the nodes showed degree of the interactions.

The correlation analysis was also performed on 35 phyla and 354 KEGG functional pathways. A total of 10 phyla strongly interacted with 64 pathways ((Spearman’s |r| > 0.7, p <0.05). A network was constructed based on 157 strong and significant correlations (84 positives and 73 negatives) between microbial phyla and KEGG functional pathways (Figure 9b). In this network, microbial phyla showed a higher degree of interactions than with the ARGs, among them Proteobacteria, Euryarchaeota, Firmicutes, Candidatus cloacimonetes, and Chloroflexi formed main large hubs.

### 3.7. Effect of treatment processes on nutrient and physio-chemical properties of pig slurry

The concentrations of main crop nutrients (N, P, K), elements (Ca, Mg, S, Na, C) and dry matter (DM) content in the control and treated samples at the end of their treatments were measured to identify if treatment resulted in a change of nutritional value for fertilisation (Table S5). The DM content decreased after storage, while it largely increased after pig slurry composting and remained almost the same in the AD system. The organic matter (OM) content increased after all the treatments. In comparison with the samples prior to treatment (control samples), the storage treatment showed enrichment of N and K, AD treatment revealed an increase in the N concentration, while composting led to a reduction in concentration of all the main nutrients. The storage also led to an increase of Na, S, and Mg elements, while reduced others. Pig slurry composting resulted in an increase of carbon with a decrease in other elements.

### 3.8. Data derived decision tree

Using the data derived from this study we generated a simple decision tree matrix (Figure 10). Each treatment was assigned a positive integer where the outcome was positive and a negative integer where the outcome was negative. The tree is based on three problems: AMR, pathogens, and Nitrogen. As the requirement for nitrogen may be either positive, in terms of maintaining the nitrogen content of the slurry or negative in terms or requiring the reduction of nitrogen to minimise pollution this was factored in two separate equations. The optimal treatment for AMR reduction was AD, for pathogen reduction were storage or composting, and for nitrogen reduction was composting but increasing nitrogen content were storage and AD. Overall, where nitrogen reduction was required, composting was the recommended treatment but where nitrogen increase was preferred then storage or AD were recommended.

**Figure 10.**
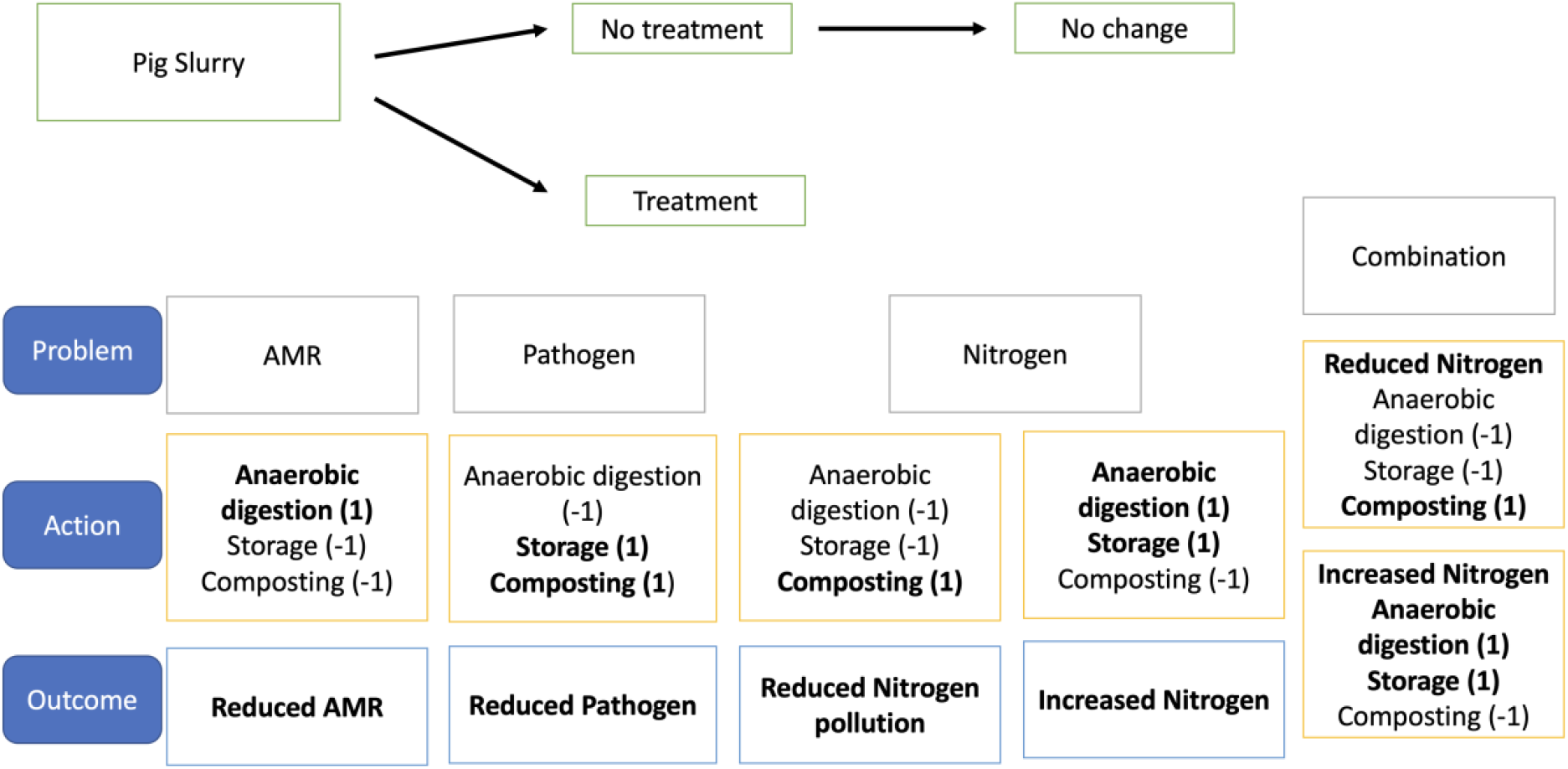
Slurry treatment decision tree. The tree is based on three problems: AMR, pathogens, and Nitrogen. Each treatment was assigned a positive integer where the outcome was positive and a negative integer where the outcome was negative.

## 4. Discussion

In this study, we analysed the effect of different mitigation treatments on human pathogen content, community composition, the resistome, and functional pathways within microbial communities in pig slurry and on slurry nutrient properties. Previous works detected major food-borne pathogens such as *Campylobacter, Salmonella,* and *Listeria* in pig slurry by culture-based techniques (Farzan et al., 2010; Mieszkin et al., 2009; Peu et al., 2006). In this work, in addition to analysis of faecal indicators, we also enumerated other critically important pathogens which pose risks to human and animal health (*Acinetobacter*, *Klebsiella*, *Pseudomonas*, and *Staphylococcus*). To the best of our knowledge the enumeration of these bacteria has not been previously reported in pig slurry under different treatments. In general, the reduction in bacterial counts was observed across all treatments, which agreed with previous findings for faecal indicators and food-borne pathogen (9, 33, 34, 35, 36). The significant decrease of faecal indicator bacteria (enterococci and *E. coli*) during storage was observed during the storage of pig slurry previously (between storage tank and the pond, and in batch studies) (Munch et al., 1987; Olsen, 1988; Peu et al., 2006). Storing waste in general reduced the abundance of enteric bacteria and could also significantly alter the compositions of enteric bacterial populations (Duriez and Topp, 2007). We detected *E. coli* in the compost samples collected only within the first 2 weeks. In comparison the *E. coli* population was stable for several weeks of storage. Most bacterial pathogens analysed were not detected at week 4 of compost except for *Enterococcus* spp.. Microbial analysis of compost at different time points in the work of McCarthy et al (2011) showed that *E. coli* decreased below the limit of detection by day 14 and remained under detection limits until day 56, while enterococci were found to day 28 and below the limit of detection by day 56 (Mc Carthy et al., 2011). In the present study the number of enumerated bacteria decreased at the beginning of AD and remained stable for the experiment duration. AD treatment has been reported to result in significant reductions of bacteria counts including coliforms (Costa et al., 2017). In general, the greater reduction of enumerated bacteria was found in stored and compost samples compared with AD samples, considering the bacterial counts in control and in the final treated products in our work. It was also noted that the levels of AMR in the bacteria tested were extremely low in all sample types, indicating that the detected potential pathogenic species were antibiotic susceptible. In the compost procedure, temperature plays an important role in the product sanitation to ensure it is generally regarded as safe to use (Malińska et al., 2014). The temperature in all compost treatment replicates was above 50°C for more than 10 consecutive days, indicating a thermophilic phase. In this phase, the thermophilic microorganisms take part in the degradation of complex compounds such as cellulose, lignin, and fats (Bernal et al., 2009). The concentration of the main nutrients varied across the different treatments. The reduction in N, P, K after pig slurry composting could make the compost product suitable to sustainable environmentally friendly agricultural land spreading. The increase in N concentration after storage and AD may cause problems with further use as fertilisers in relation to water pollution and climate change policy implementation in the future.

### 4.1. Effect of treatment on microbial communities

Using manure as soil fertilisers could introduce both beneficial and pathogenic microorganisms into soils. Previous studies on pig manure treatment mainly focused on analysing the microbial communities under one treatment, such as different storage conditions (Lim et al., 2018) or using different raw material contents, or at different stages of AD or composting procedures (Demirel and Scherer, 2008; Liu et al., 2020; Partanen et al., 2010; Ros et al., 2017). Here, we presented a first study investigating the dynamics of microbial communities during different treatments (storage, composting, and AD) on the same pig slurry. A major expectation of this study was to compare the microbial response across all treatments to address the efficiency and molecular basis behind the result of each treatment process. The pig slurry treatments analysed in our work showed the effects on microbial community composition differed by treatment, indicating that each treatment created distinct microbial communities in slurry. In common with previous reports, at the phylum level, we detected the predominance of Firmicutes, Bacteroidetes, and Proteobacteria, which were the most abundant phyla in the gut microbiome of pigs (Tang et al., 2020; X. Wang et al., 2019). The dominance of these phyla detected during storage was also found in the study of Kumar et al.,(Kumar et al., 2020). The reduction in relative abundances of Firmicutes and the increase of Bacteroidetes were also consistent with the results of other studies of pig slurry microbiota of pig slurry during storage and chicken guts at different growing time (Kumar et al., 2020; Mohd Shaufi et al., 2015). *Prevotella*, *Bacteroides*, and *Lactobacillus* were dominant in microbial communities of stored pig slurry at the genus level. These bacteria were also reported as the dominant taxa in storage of pig slurry previously (9, 27, 39).

We observed the dominance of bacterial phyla Firmicutes, Bacteroidetes, Proteobacteria, and the archaea phylum Euryachaeota during the AD treatment. These phyla were commonly detected during AD in previous works (Di Maria and Barratta, 2015; Ros et al., 2017; Stolze et al., 2015). Moreover, the most abundant phyla detected in our work was Firmicutes which was also increased in the AD process. Firmicutes were reported as the dominant bacteria in the mesophilic reactor at 37°C (Zamanzadeh et al., 2017). This phylum consists of bacteria involved in the degradation of various volatile fatty acids detected in AD and activated sludge systems (Garcia-Peña et al., 2011).

The increase of Firmicutes and the decrease in relative abundance of Bacteroidetes, Proteobacteria, and Actinobacteria detected here was also reported by different authors (Di Maria and Barratta, 2015; Ros et al., 2017; Stolze et al., 2015). Indeed, Ros et al 2017 (Ros et al., 2017) indicated this change in abundance of these phyla when studying mesophilic AD of pig slurry and fruit and vegetable waste. Stolze et al., 2015 (Stolze et al., 2015) also found the trend when analysing the operation of an agricultural biogas reactor under different conditions, as was observed in the work of Nelson et al, 2011 (Nelson et al., 2011) which studied anaerobic digesters using a meta-analysis method. The Euryarchaeota was the most abundant Archaea, which was also frequently identified in other AD studies (Leclerc et al., 2004; Pampillón-González et al., 2017; Rabii et al., 2019). The microbial community composition of the AD samples at the genus level was dominanted by 3 bacteria genera *Syntrophomonas*, *Hungateiclostridium*, *Clostridium* (belonging to Firmicutes), and 2 Archaea genera *Methanoculleus*, and *Methanobacterium* (belonging to Euryarchaeota). These bacterial phyla participate in syntrophic metabolism, substrate hydrolysis, and fermentation (Bosshard et al., 2002; Chen et al., n.d.; Vanwonterghem et al., 2014). *Clostridium* is known to dominate the hydrolytic and acidogenic stages of AD (Fontana et al., 2016). It is important to address the presence of any potential pathogenic species in this genus in AD products, which could pose risks to the agricultural environment, humans, and animals. Both archaea genera are hydrogenotrophic methanogens (Ros et al., 2017).

Similar to previous studies (Jiang et al., 2020; Liu et al., 2020; Zhang et al., 2018), the 4 most abundant phyla during composting were Proteobacteria, Bacteroidetes, Firmicutes, and Actinobacteria. Proteobacteria was the most dominant bacterial phylum. The bacterial community composition underwent a succession from Firmicutes dominance before composting to an abundance of Proteobacteria and Bacteroidetes during composting within the first two weeks. Proteobacteria, Bacteroidetes, and Actinobacteria have been reported to play important roles in the degradation of organic matter in composting (Awasthi et al., 2017), while Firmicutes participate in the decomposition of lignin, cellulose, and hemicellulose (PANDEY et al., 2013). The decrease in the relative abundance of Firmicutes along with the increase of Proteobacteria and Bacteroidetes in our work was in line with previous studies (Jiang et al., 2019; Li et al., 2020). Firmicutes can grow at high temperatures, thus these bacterial phyla may dominate the beginning of composting. Proteobacteria and Bacteroidetes can likely exist at lower temperatures in the late composting stages. We observed a dominance of the *Pseudomonas* genus in compost treatment samples. *Pseudomonas* was reported to be highly productive at different composting stages. These bacteria dissolve minerals, produce nutrients, and play an important role in lipid degradation. Thus, they contribute to the quality of the compost product as soil fertilisers.

The LEfSE analysis identified the microbial fingerprints in our data of genus abundances for each treatment sample group. These fingerprints can describe the treatment-specific taxa by comparing the taxa abundances between different sample types (28). The LEfSE results indicated that the microbial taxa present could be used to differentiate treatment samples from each other and from control groups. These also determined different responses of microbial communities to different conditions between control and all treatment groups, establishing the unique characteristic microbiota in each sample group. These highlighted the important role of treatment-specific taxa as major factors driving the microbiome functions in each treatment type. The LEfSE analysis provided information for a better understanding of microbial communities and can be used to identify novel metagenomic biomarkers (28).

### 4.2. Microbial diversity

The PCoA analysis based on the Bray-Curtis dissimilarity matrices showed 4 distinct clusters formed by the 4 sample groups, indicating the significant difference in the microbial community structures across sample groups. This also confirmed the individual microbial profile of each treatment, which aligned with the relative abundance of microbial taxa. Similarly, the overlap between 2 clusters of the storage and control samples in the PCoA plot confirmed the prevalent order of microbial phyla within the samples. These results indicated that the sample classification can be established based on the microbial community in the sample and that the treatments each alter the microbial content of the slurry in a unique manner, except storage.

The alpha diversity (Chao1 and Shannon indexes) can be used to identify the variation of microbial diversity between samples. The total microbial biomass examined in the Chao 1 index for microbial richness was significantly increased after storage and reduced after compost and AD. The relatively rich nutrient environment in the storage samples would be responsible for the increase of microbial richness by promoting copiotrophic microorganisms (Medina-Sauza et al., 2019). A similar variation was also found with the Shannon diversity across all sample groups. Overall, the Shannon indexes also reduced in AD and compost treatments but increased in storage. The decrease of Shannon diversity was also reported by Wan et al 2021 (Wan et al., 2021) when analysing the microbial composition during composting, and Zealand et al studying anaerobic co-digestion of dairy manure with rice straw (Zealand et al., 2018). However, our study demonstrates the relative variations across the same input sample when treated in different manners. The Shannon diversity index has been used to assess community diversity. The difference in alpha diversity across different sample groups may be due to the availability of nutrient contents (Shehata et al., 2021). Indeed, the increase of Shannon index in the storage is likely due to the rich nutrient contents during the treatment, while the lowest alpha diversity indexes in compost samples could relate to the lowest nutrient contents in this process.

### 4.3. Resistome analysis

Our study was the first to analyse and compare the resistome in pig slurry across storage, composting and AD treatments. The profiles of ARGs and MGEs were different between all sample groups. This result was confirmed by separated clusters in the PCoA plot. The storage samples displayed the most similar profile to control in comparison with other treatments, which was also indicated by the overlap between two clusters. The relative abundance of both ARGs and MGEs was lowest in AD treatment, indicating the highest efficiency in removing antibiotic resistance. These also resulted in the low richness (Chao1) of ARGs and MGEs in AD sample groups comparing with control and other sample groups. However, the Shannon diversity index of ARGs was found at the highest value in AD among all samples, indicating a high diversity of ARGs and MGEs present. The increase in total ARGs during pig slurry composting was reported previously. Cao et al identified an increase by 0.19-1.62 logs of the relative abundance of total ARGs after composting (Cao et al., 2020). The relative abundance of ARGs started to increase at week 2 of composting compared with control samples. This decreased in weeks 6 and 8 but remained at a higher level than the other treatments. Similar results were also reported by other authors in studying pig slurry and sewage sludge composting (Cao et al., 2020; Su et al., 2015; Wang et al., 2015). However, few other studies showed any decrease in ARGs in pig slurry composting (Chen et al., 2007; Selvam et al., 2012). The effect of compost procedures on ARG removal efficiency has differed among many studies (Chen et al., 2007; Selvam et al., 2012; Wang et al., 2015; Zhang et al., 2017). The fate of different kinds of ARGs varied and the abundances of some ARGs increased while others decreased during/after composting (Zhang et al., 2017). The behaviour of ARGs in the same class was also different, for instance, *tetM*, *tetO*, *tetQ*, and *tetW* were decreased while *tetA*, *tetC*, *tetG* and *tetL* increased after pig slurry composting (Wang et al., 2015). The increase of aminoglycoside and sulfonamide resistance here is also in agreement with other studies (Cao et al., 2020; Wang et al., 2015). It is important to note that, the compost in this study was kept at 50°C for over 10 days, but higher temperatures were applied by other authors, which may alter the ARG removal efficiency. The efficiency of compost in removing ARGs was not always satisfactory; it strongly depends on the properties of the compost mixture and the control conditions. The core resistome of pig slurry among all sample groups, with a high abundance of tetracycline and aminoglycoside resistance genes, was consistent with the core antibiotic resistome of gut microbiota in industrialised feedlot pigs and laboratory pigs (Looft et al., 2012; C. Wang et al., 2019).

The relationship between ARGs and MGEs was visualised in the network based on Spearman’s correlation analysis. The high degree of positive correlations with various ARGs was found for the intI1 integron, which is a proxy for anthropogenic gene pollution (16). A high number of genes within the tetracycline, aminoglycoside, and sulfonamide resistance gene groups positively interacted with MGEs in all sample. This result indicates the strong dissemination of these genes via horizontal gene transfer. This was also reported in previous works analysing pig slurry (Cao et al., 2020; Wang et al., 2021). The integrons and transposons formed the main large hubs in the network, determining their essential role in ARG dissemination.

The strong interaction between ARGs and microbial phyla was found in the correlation analysis. Most ARGs showed multiple interactions with different phyla, indicating a range of potential ARG bacterial hosts. This was consistent with previous findings (Qian et al., 2021; Wang et al., 2021). The main hubs of the network were formed by microbial phyla, showing their essential role in network formations. Proteobacteria and Bacteroidetes were the primary ARGs carriers, identified through the most positive interaction with ARGs. Bacteroidetes were strongly correlated to *tet* genes, which was also previously reported, and we know that many members of the Bacteroidetes phyla contain *tet* genes on their chromosomes (Wang et al., 2017). Actinobacteria were identified as carriers of *tetW* and *bla*_CTX-M-6_. Firmicutes were found to be important in ARG dissemination (Song et al., 2017). This phylum showed positive interactions with aminoglycoside resistance genes, with negative interactions with others. These bacterial phyla were also reported as the ARG hosts in soil microbiota (Qian et al., 2021). Euryachaeota and Candidatus Cloacimonetes held a high degree of negative interactions with ARGs. The KEGG pathway analysis identified the representative functional pathways for each sample group as well as the core pathways among all samples. The ABC transporters was the most abundant in our work, which was previously reported among the high abundance species (Campanaro et al., 2016). The core pathways contain almost all annotated pathways, the alpha diversity of functional pathways was significantly higher after all treatments. This result showed that many pathways were common among all sample groups, but their relative abundance differed significantly, indicating the difference in their availability for utilisation across all samples. This could also indicate a high metabolism level in all the treatments. PCoA analysis confirmed the variation of the difference in profile and composition of detected pathways in all sample groups by 4 distinct clusters. The dominance of methane metabolic pathways in the AD treatment was consistent with previous findings (Guo et al., 2015; Ma et al., 2021). This is directly linked to the high abundance of methanogenic archaea in the microbial community. The network analysis also identified the strong interactions of Proteobacteria, Euryarchaeota, Firmicutes, Candidatus Cloacimonetes, and Chloroflex with KEGG pathways.

## 5. Conclusions

This is the first study to provide a direct comparison of different treatments on the same pig slurry microbiome and resistome. Our results determined the effectiveness of all three treatments (storage, composting, and AD) on reducing/removing potential pathogens by culture-based methods. Our work also revealed the distinct response of the pig slurry microbiome under different conditions. Indeed, storage is considered the simplest technique to treat pig slurry in comparison to composting and AD. However, composting and AD showed better capacities to decrease the diversity of microbial communities, especially AD which also showed the best efficiency at reducing the microbial load, ARGs and MGEs in pig slurry. The link between microbial composition and resistome is well demonstrated in our study via the compost samples, where the change in microbial composition resulting in Proteobacteria and Bacteroidetes dominating, is mirrored by a large increase in the relative abundance of ARGs and MGEs. This indicates that the changes in microbial population composition correlate with specific ARG and MGE changes in the populations. Such data is only possible due to the extensive sequencing and data analysis performed in this study. Similar microbial changes have been observed in previous studies demonstrating the reliability of these data. Although the treatments in our work were designed on small scale and more data should be obtained for the nutrient quality, the outcomes can be used to design an optimal low-cost treatment adapted to actual conditions on different pig farms. The most suitable treatment process will depend on scenarios such as geographical area, the desired quality of the final product, the farm size, and the associated economic cost. The results of our work can provide the tools for farmers to determine appropriate solutions and make the decision for direct treatments on the farm. Our results showed that each of the treatment methods had advantages and disadvantages, depending on the parameter measured e.g., reduction in ARG or MGE content or microbial diversity or pathogens. However, the implications of the changes within the microbiome are not yet known. Our data derived decision tree provides a structure for the determination of optimal strategies for slurry treatment. While this is a simple model of decision additional factors, such as cost, or time may be added to determine the optimal recommendation. The most important component of this tree is that the data is comparable as all treatments were performed on the same initial slurry samples, thus the only changing component was the treatment. Future decision trees and models may be generated from these data to model different input data such as higher or lower pathogen content or different AMR gene or nutrient content.

Other aspects related to the farm scale, desired products, and the cost for treatment installation and performance control should be further analysed. For future work, a combination of variable methods in the treatment process should be examined to select the best available technologies for the best utilisation approach in a given region.

## Supporting information

Supplemental information all files

## Author Contributions

**TTD and FW:** Design and/or interpretation of the reported experiments, Acquisition and/or analysis of data, Drafting and revising the manuscript, Administrative, technical or supervisory support. **SN**: Acquisition and/or analysis of data, Drafting and revising the manuscript. **NH**: Acquisition and/or analysis of data. **VOF**: Administrative, technical, or supervisory support. **FB and CB:** Drafting and revising the manuscript, Administrative, technical, or supervisory support.

## Acknowledgments

The authors would like to thank the Irish pig farm for supporting our sample collection. We also thank the farm and technical staff at Teagasc.

## Findings

This work was funded by Irish Health Research Board, in the frame of the JPI-EC-AMR Joint Transnational Call (JPIAMR), JPI-EC-AMR JTC 2017, project INART – “Intervention of antibiotic resistance transfer into the food chain” to FB and FW.

## Appendix A. Supplementary Material

The following are Supplementary data to this manuscript:

- Supplementary figures and tables
- Datasets S1 to S5

