## Supplemental information all files for "Data based slurry treatment decision tree to minimise antibiotic resistance and pathogen transfer while maximising nutrient recycling"

**The supplementary materials include:**

- Legends for Datasets S1 to S5
- Figures S1 to S8(not allowed for Brief Reports)
- Tables S1 to S5 (not allowed for Brief Reports)

**Data legends:**

**Dataset S1.** Relative abundances (%) of microbial phyla

**Dataset S2.** Relative abundances of resistance gene classes, ARGs, and, MGEs

**Dataset S3.** Correlation analysis of ARGs and MGEs

**Dataset S4.** Correlation analysis of bacterial phyla with ARGs and KEGG functional pathways

**Dataset S5.** HT-qPCR gene relative abundances

**Figures:**

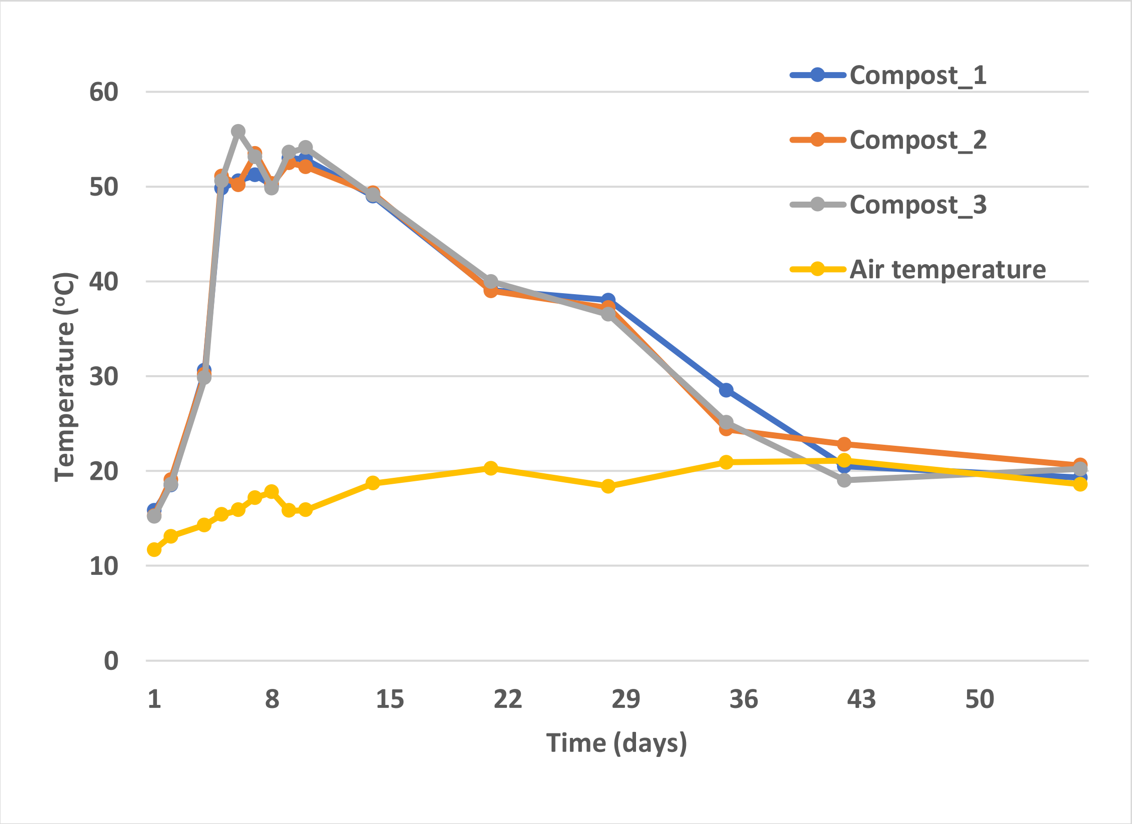


**Figure S1.** Temperature recorded during the composting process of pig slurry.

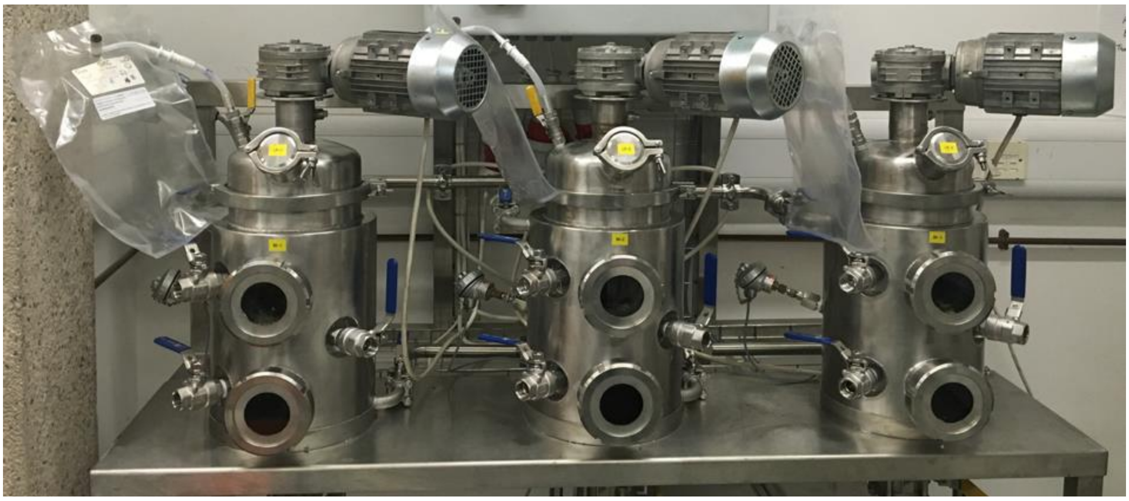

**Figure S2.** Triplicate 10 L continuously stirred tank reactors (CSTRs).

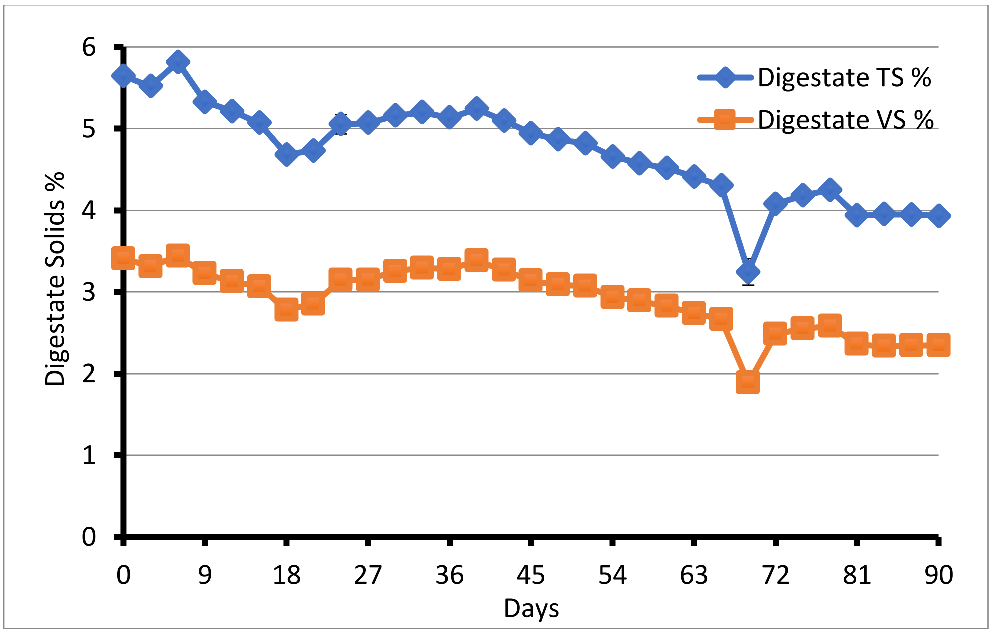


**Figure S3.** Total (TS) and volatile solids (VS) in the digestate over time.

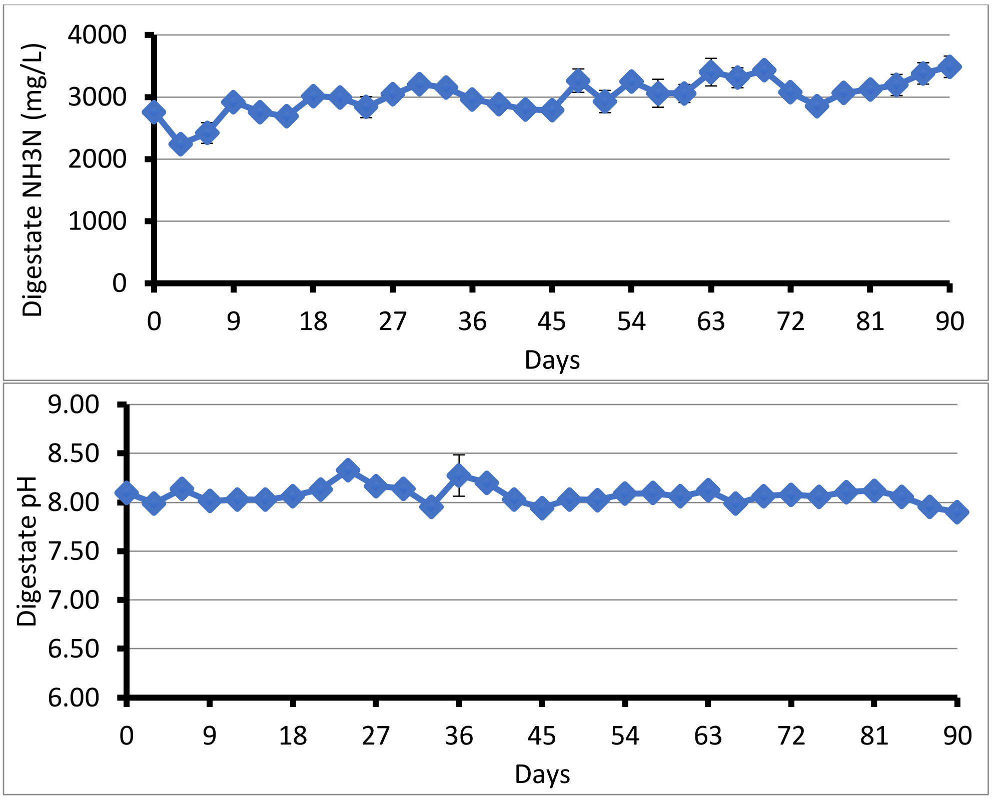


**Figure S4.** $NH_3$-N concentratiol (mg/L) and pH in the digestate over time.

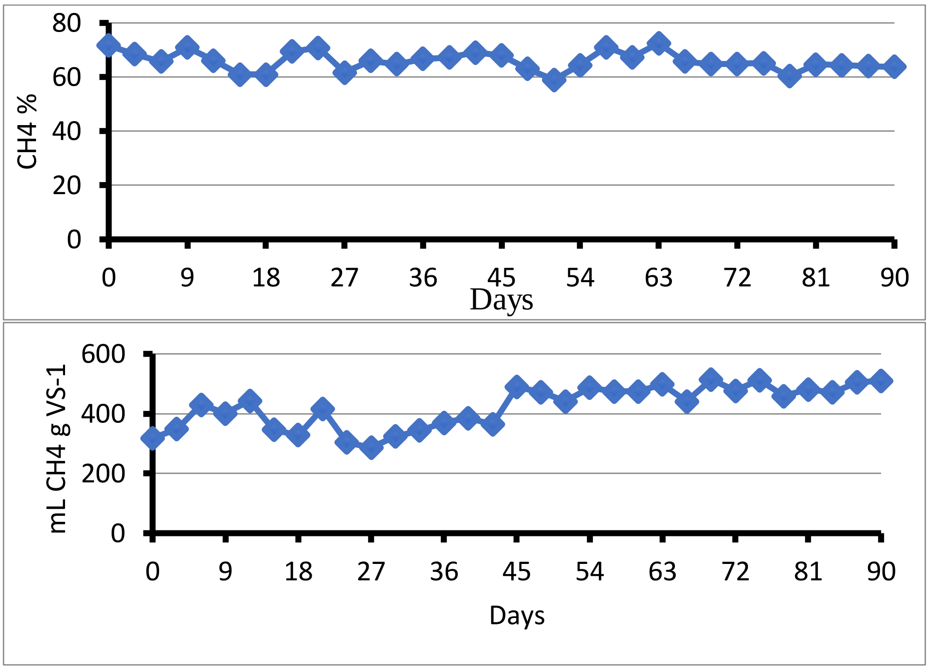


**Figure S5.** $CH_4$ production in the digestate over time.

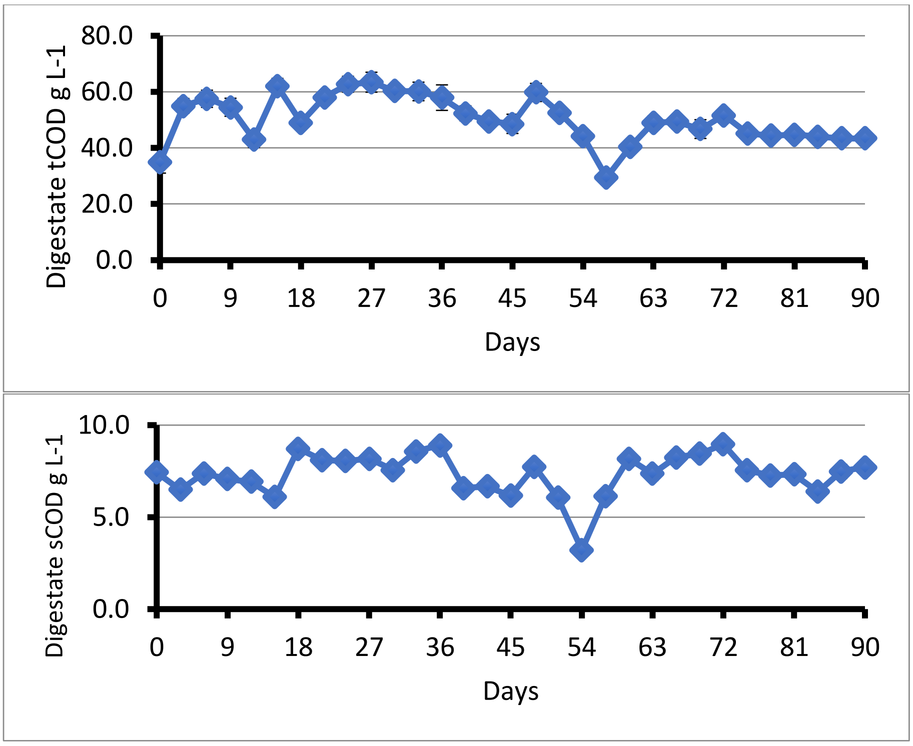


**Figure S6.** Total COD in the digestate over time.

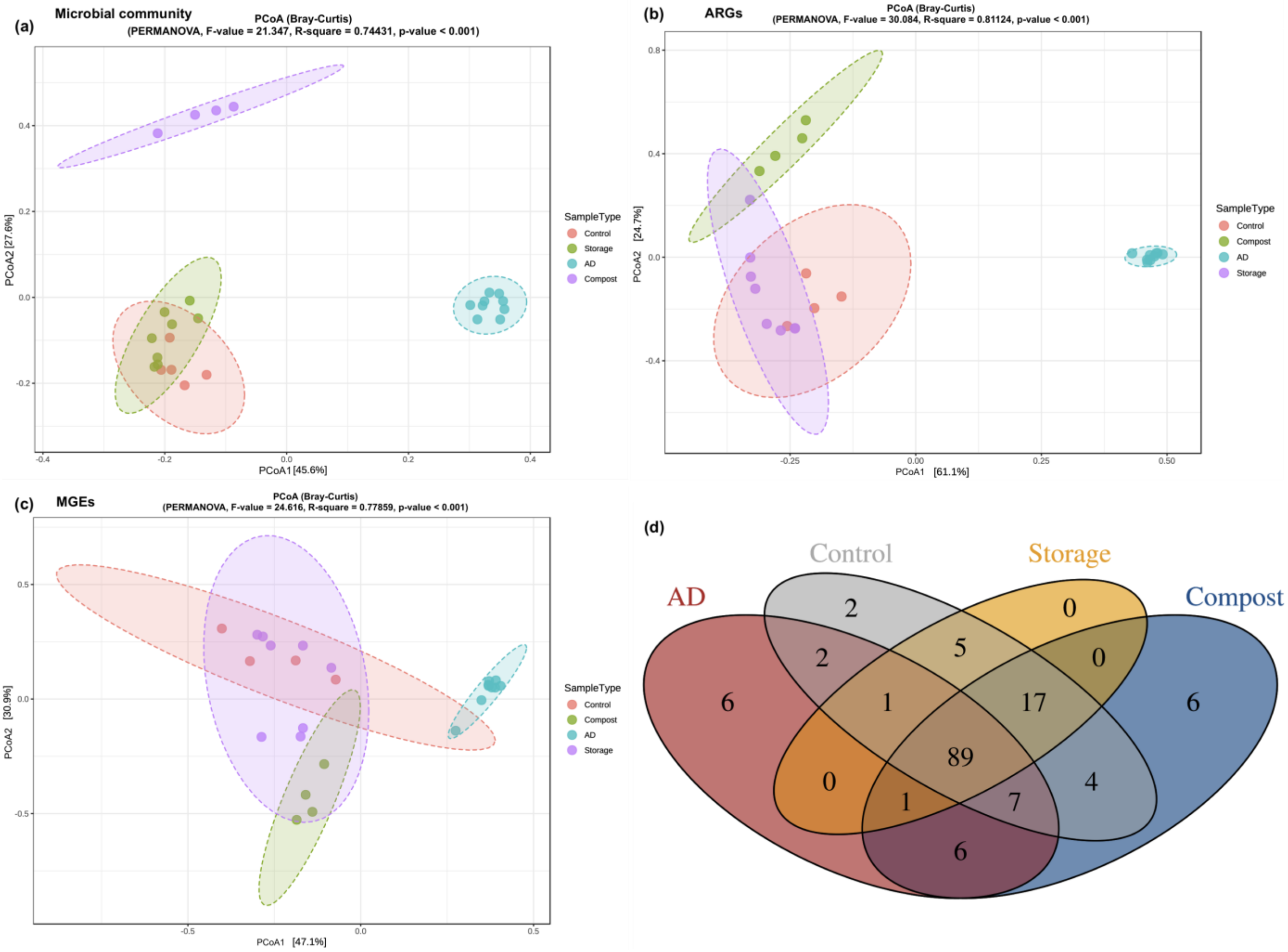


**Figure S7.** The composition of microbiome and resistome in pig slurry. Principal coordinates analysis (PCoA) based on the Bray-Curtis dissimilarity matrices of microbial community compositions in all sample (a), of ARGs (b), and MGEs (c). Core resistome in pig slurry across all sample groups (d). Venn diagram presenting the number of detected genes among study samples and their intersection.

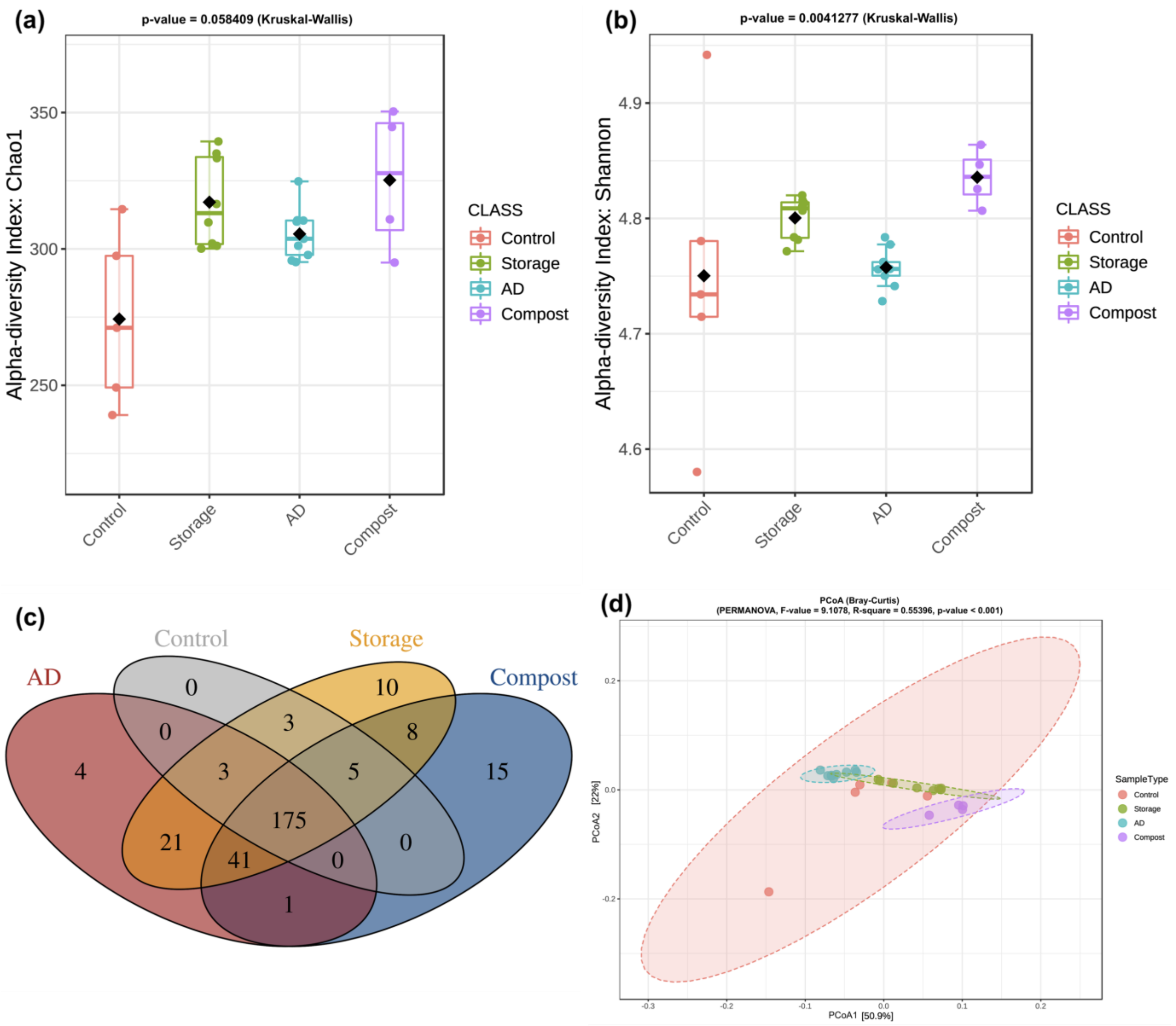


**Figure S8.** Alpha diversity of KEGG functional pathways (a, b). The core pathways in pig slurry across all sample groups (c). Venn diagram presenting the number of detected pathways among study samples and their intersection. Principal coordinates analysis (PCoA) of KEGG functional pathways in all sample based on the Bray-Curtis dissimilarity matrices (d).

**Tables:**

**Table S1.** Average physicochemical data of pig slurry prior AD and mixed feedstock throughout trial.

| | TS | VS | NH3-N | tCOD | sCOD | pH |
|---|---|---|---|---|---|---|
| | % | % | mg/L | g/L | g/L | |
| Pig slurry | 3.14±0.13 | 2.01±0.12 | 2963±27 | 80±5.8 | 36±0.94 | 6.31±0.05 |
| Feedstock | 9.96±0.31 | 8.46±0.24 | 2269±32 | 221±9.3 | 31±2.1 | 5.54±0.02 |

**Table S2.** Conditions and selective agars with/without antibiotics (at EUCAST breakpoint concentrations) used to enumerate potential pathogens.

| **Bacteria** | **Selective Agar + antibiotic** | **Selecting for** | **Incubation conditions** |
|---|---|---|---|
| *E.coli* | EMB (Oxoid) | *E. coli* | 37°C for 24 hours |
| *E.coli* | EMB + Cefotaxime (4 mg/L) | ESBL | 37°C for 24 hours |
| *E.coli* | EMB + Imipenem (16 mg/L) | Carbapenem resistance | 37°C for 24 hours |
| *Klebsiella spp.* | Simmon citrate agar + 1% Inositol (Oxoid) | *Klebsiella spp.* | 37°C for 24 hours |
| *Klebsiella spp.* | Simmon citrate agar + 1% Inositol + Cefotaxime (4 mg/L) | ESBL | 37°C for 24 hours |
| *Klebsiella spp.* | Simmon citrate agar + 1% Inositol + Imipenem (16 mg/L) | Carbapenem resistance | 37°C for 24 hours |
| *Acinetobacter spp.* | Leeds Acinetobacter Medium (HIMEDIA) | *Acinetobacter spp.* | 37°C for 24 hours |
| *Acinetobacter spp.* | Leeds Acinetobacter Medium + Imipenem (16 mg/L) | Carbapenem resistance | 37°C for 24 hours |
| *Pseudomonas spp.* | Pseudomonas Isolation Agar (Oxoid) | *Pseudomonas spp.* | 37°C for 24 hours |
| *Pseudomonas spp* | Pseudomonas Isolation Agar + Imipenem (16 mg/L) | Carbapenem resistance | 37°C for 24 hours |
| *Staphylococcus* spp. | Mannitol salt agar (Oxoid) | *Staphylococcus* spp. | 37°C for 24-48 hours |
| *Staphylococcus* spp. | Mannitol salt agar (Oxoid) + Oxacillin (4 mg/L) | Methicillin resistance | 37°C for 24-48 hours |
| Enterococci | Enterococcus selective agar (Sigma-Merck) | Enterococci | 37°C for 24-48 hours |
| Enterococci | Enterococcus selective agar + Vancomycin (32 mg/L) | Vancomycin resistance | 37°C for 24-48 hours |

**Table S3.** Unclassified reads and taxonomic classifications from classified reads in Kaiju results. F0 = control samples, CP = compost samples, AD = anaerobic digestion samples, PS = storage samples.

| **Sample ID** | **Unclassified reads (%)** | **Bacteria of classified reads (%)** | **Archaea of classified reads (%)** | **Virus and others (%)** |
|---|---|---|---|---|
| F0-1 | 46.59 | 97 | 1 | 2 |
| F0-2 | 45.52 | 96 | 2 | 2 |
| CP-W2 | 26.62 | 99 | 0.4 | 0.6 |
| CP-W4 | 23.3 | 99 | 0.5 | 0.5 |
| CP-W6 | 21.71 | 99 | 0.2 | 0.8 |

| | | | | |
|---|---|---|---|---|
| CP-W8 | 21.56 | 99 | 0.2 | 0.8 |
| F0-3 | 48.79 | 96 | 1 | 3 |
| AD-W1 | 32.47 | 91 | 8 | 1 |
| AD-W2 | 34.9 | 96 | 4 | 0 |
| AD-W4 | 33.05 | 94 | 6 | 0 |
| AD-W6 | 32.35 | 93 | 7 | 0 |
| AD-W8 | 33.32 | 94 | 6 | 0 |
| AD-W10 | 33.85 | 94 | 6 | 0 |
| AD-W12 | 33.28 | 92 | 8 | 0 |
| AD-W14 | 33.77 | 90 | 10 | 0 |
| AD-W16 | 35.67 | 95 | 5 | 0 |
| F0-4 | 59.66 | 92 | 3 | 5 |
| F0-5 | 55.95 | 94 | 1 | 5 |
| PS-W2 | 51.59 | 96 | 0.8 | 3.2 |
| PS-W4 | 50.79 | 96 | 0.9 | 3.1 |
| PS-W6 | 54.88 | 95 | 1 | 4 |
| PS-W8 | 55.87 | 95 | 0.8 | 4.2 |
| PS-W10 | 53.87 | 95 | 0.7 | 4.3 |
| PS-W12 | 56.5 | 95 | 0.9 | 4.1 |
| PS-W14 | 53.52 | 95 | 0.8 | 4.2 |
| PS-W16 | 52.92 | 95 | 0.7 | 4.3 |

**Table S4**. Share ARGs and MGEs among resistome of control and treatment sample groups. Core = shared across all samples.

| **Gene** | **Gene Type** | **Resistance phenotype /MGEs** |
|---|---|---|
| *strB* | core | Aminoglycoside |
| *aphA3_2* | core | Aminoglycoside |
| *aphA3_1* | core | Aminoglycoside |
| *ampC_6* | core | Aminoglycoside |
| *aadE* | core | Aminoglycoside |
| *aadD* | core | Aminoglycoside |
| *aadA9_2* | core | Aminoglycoside |
| *aadA5_1* | core | Aminoglycoside |
| *aadA2_3* | core | Aminoglycoside |
| *aadA_2* | core | Aminoglycoside |
| *aadA_1* | core | Aminoglycoside |
| *aadA1* | core | Aminoglycoside |
| *aac6Ib_3* | core | Aminoglycoside |
| *aac6Ib_2* | core | Aminoglycoside |
| *aac6Ib_1* | core | Aminoglycoside |
| *aac3VI* | core | Aminoglycoside |
| *cphA_1* | core | Beta-lactam |
| $bla_{SHV\_1}$ | core | Beta-lactam |
| $bla_{OXY}$ | core | Beta-lactam |
| $bla_{OXA-48}$ | core | Beta-lactam |
| $bla_{OCH}$ | core | Beta-lactam |
| $bla_{NDM}$ | core | Beta-lactam |
| $bla_{CTX-M\_4}$ | core | Beta-lactam |

| | | |
|---|---|---|
| *bla*$_{CTX\text{-}M\_2}$ | core | Beta-lactam |
| *bla*$_{CTX\text{-}M\_1}$ | core | Beta-lactam |
| *bla*$_{CTX\text{-}M}$ | core | Beta-lactam |
| *bla*$_{CMY2}$ | core | Beta-lactam |
| *ampC_2* | core | Beta-lactam |
| orf37IS26 | core | Insertional sequence |
| IS613 | core | Insertional sequence |
| IS1133 | core | Insertional sequence |
| IS1111 | core | Insertional sequence |
| intI3_2 | core | Integron |
| intI2_2 | core | Integron |
| intI1_4 | core | Integron |
| intI1_3 | core | Integron |
| intI1_2 | core | Integron |
| intI1_1 | core | Integron |
| *acrA_1* | core | MDR |
| *qacFH* | core | MDR_mobile |
| *mphA_1* | core | MLSB |
| *mefA* | core | MLSB |
| *ermB_1* | core | MLSB |
| *ermA* | core | MLSB |
| *cmxA* | core | Phenicol |
| *repA* | core | Plasmid_associated |
| IncW_*trwAB* | core | Plasmid_associated |
| IncQ_oriT | core | Plasmid_associated |
| IncP_oriT | core | Plasmid_associated |
| IncN_rep | core | Plasmid_associated |
| *mcr1* | core | Polymyxin |
| *qepA* | core | Quinolone |
| *sul3_1* | core | Sulfonamide |
| *sul2_2* | core | Sulfonamide |
| *sul2_1* | core | Sulfonamide |
| *sul1_2* | core | Sulfonamide |
| *sul1_1* | core | Sulfonamide |
| *tetW* | core | Tetracycline |
| *tetR_3* | core | Tetracycline |
| *tetR* | core | Tetracycline |
| *tetO_2* | core | Tetracycline |
| *tetO_1* | core | Tetracycline |
| *tetM_2* | core | Tetracycline |
| *tetM_1* | core | Tetracycline |
| *tetM* | core | Tetracycline |
| *tetH* | core | Tetracycline |
| *tetG_2* | core | Tetracycline |
| *tetG_1* | core | Tetracycline |
| *tetG* | core | Tetracycline |
| *tetD_3* | core | Tetracycline |
| *tetD* | core | Tetracycline |
| *tetAP* | core | Tetracycline |
| *tetAB_1* | core | Tetracycline |

| | | |
|---|---|---|
| *tetA_2* | core | Tetracycline |
| Tp614 | core | Transposon |
| tnpA_7 | core | Transposon |
| tnpA_6 | core | Transposon |
| tnpA_5 | core | Transposon |
| tnpA_3 | core | Transposon |
| tnpA_2 | core | Transposon |
| tnpA_1 | core | Transposon |
| *dfrA1_1* | core | Trimethoprim |
| *dfrA1* | core | Trimethoprim |
| *vanHD* | core | Vancomycin |
| *vanHB* | core | Vancomycin |
| *vanC_3* | core | Vancomycin |
| *vanB_3* | core | Vancomycin |
| *vanB_1* | core | Vancomycin |
| *vanA* | core | Vancomycin |

**Table S5**. Dry matter, organic matter, and major nutrient contents in the control samples (raw slurry) and the treatment products at the end of the treatment process.

| Sample type | DM (% of WT) | OM (% of DM) | Total N (g/kg) | Total P (g/kg) | Total K (g/kg) | Total C (% of DM) | Ca (g/kg) | Na (g/kg) | S (g/kg) | Mg (g/kg) |
|---|---|---|---|---|---|---|---|---|---|---|
| **Control** | 4.18±0.18 | 54.94±3.77 | 42.03±0.58 | 12.41±0.3 | 31.97±0.38 | 41.1±0.12 | 23.1±0.53 | 6.93±0.04 | 8.91±0.11 | 7.81±0.22 |
| **Storage** | 3.14±0.04 | 65.65±1.46 | 47.4±0.36 | 11.83±0.19 | 51.57±0.34 | 38.27±0.33 | 18.89±0.07 | 12±0.06 | 10.73±0.31 | 7.38±0.2 |
| **Compost** | 22.33±0.18 | 79.06±1.46 | 17.8±1,09 | 6.61±0.71 | 14.08±2.31 | 44.2±0.21 | 11.2±0.44 | 2.4±0.3 | 4.55±0.29 | 2.93±0.27 |
| **AD** | 4.73±0.11 | 61.5%±0.33 | 49±2.08 | 9.27±0.43 | 22.67±0.32 | 29.17±0.15 | 11.14±0.57 | 6.81±0.26 | 3.32±0.37 | 2.84±0.16 |
